# Temperature sensitive Mutant Proteome Profiling: a novel tool for the characterization of the global impact of missense mutations on the proteome

**DOI:** 10.1101/2019.12.30.891267

**Authors:** Sarah A. Peck Justice, Guihong Qi, H. R. Sagara Wijeratne, José F. Victorino, Ed R. Simpson, Aruna B. Wijeratne, Amber L. Mosley

## Abstract

Temperature sensitive (TS) mutants have been foundational in the characterization of essential genes. However, a high-throughput workflow for characterization of biophysical changes in TS mutants is lacking. <u>Te</u>mperature sensitive <u>M</u>utant <u>P</u>roteome <u>P</u>rofiling (TeMPP) is a novel application of mass spectrometry (MS) based thermal proteome profiling (TPP) to characterize effects of missense mutations on protein stability and PPIs. This study characterizes missense mutations in two different subunits of the 26S proteasome on the thermal stability of the proteome at large, revealing distinct mechanistic details that were not obtained using only steady-state transcriptome and proteome analyses. TeMPP is a precise approach to measure changes in missense mutant containing proteomes without the requirement for large amounts of starting material, specific antibodies against proteins of interest, and/or genetic manipulation of the biological system. Overall, TeMPP provides unique mechanistic insights into missense mutation dysfunction and connection of genotype to phenotype in a rapid, non-biased fashion.

## INTRODUCTION

The ability to systematically assess the function of individual genes and gene products is critical for gaining a full picture of how the cell works (Boone, 2014). Much of our advancement in knowledge on biological processes and the underlying mechanisms is owed to the use of genetic methodologies that perturb normal gene function (discussed in (Nurse & Hayles, 2019; Winston & Koshland, 2016)). One such molecular biology tool that is particularly useful for studying essential gene function are conditional mutants. These include temperature sensitive (TS) mutants and mutants which show phenotypic defects under conditions such as starvation. have been instrumental for decades in the study of many essential biological processes including but not limited to: characterization of the cell cycle (Hartwell, 1971; Hartwell, Culotti, & Reid, 1970; Hartwell, Mortimer, Culotti, & Culotti, 1973; Lee & Nurse, 1987; Nurse & Bissett, 1981; Nurse, Thuriaux, & Nasmyth, 1976), RNA polymerase II transcription (Cormack & Struhl, 1992; Gross, Grossman, Liebke, Walter, & Burgess, 1984; Hartzog, Wada, Handa, & Winston, 1998; Holstege et al., 1998; Nonet, Scafe, Sexton, & Young, 1987; Takagi et al., 2006; Thompson & Young, 1995; Winston, Chaleff, Valent, & Fink, 1984), and autophagy (Kametaka, Okano, Ohsumi, & Ohsumi, 1998; Shirahama, Noda, & Ohsumi, 1997; Takeshige, Baba, Tsuboi, Noda, & Ohsumi, 1992; Thumm et al., 1994; Tsukada & Ohsumi, 1993), amongst others. TS mutants have distinct advantages relative to other conditional mutants in that they provide the ability to inactivate genes without having to change the transcriptional context of the gene, add chemicals, or change media (Kofoed et al., 2015). Furthermore, by simply altering the temperature at which cells are grown, TS mutant proteins can be studied in a context in which they retain function (permissive temperature), lose function (nonpermissive temperature), or have partial function (semi-permissive temperature) (Ben-Aroya et al., 2008; Sugaya, 2018).

As early as the 1960s, TS mutants have been differentiated by the mechanism of dysfunction and broken into two broad categories: temperature-sensitive folding (TSF) and thermolabile (TL) (Lovato, Adams, Baker, & Cripps, 2009; Sadler & Novick, 1965). TSF mutants only show a TS phenotype if cells are grown at restrictive temperatures during protein synthesis; if shifted once the protein is folded, no phenotype is observed. It has been postulated that these mutants are stabilized via integration into protein complexes, and therefore their interactions with other subunits stabilize them even after a temperature shift (Edgar & Lielausis, 1964; Gordon & King, 1994). TL mutants, on the other hand, display a phenotype immediately when incubated at restrictive temperatures. In this case, it is often thought that the mutant phenotype is a result of protein destabilization and/or loss of protein function. This could be occurring through multiple mechanisms including decreased ability to interact with other proteins or decreased melting temperature from the loss of amino acids crucial to proper protein folding (Lovato et al., 2009).

Despite the long-term use of TS mutants, much is still left to discover in terms of the properties and mechanisms leading to their temperature sensitivity. Many of the forward genetics screening processes that produce TS mutants result in multiple mutations which can make determination of which mutations are causative and what mechanism is underlying the altered phenotype difficult. The ability to characterize the effects of TS mutants on the cell would provide immense benefit for the use of these mutants to study essential biological pathways in various model systems. Predictive software has been developed to generate TS mutants that standard forward genetics approaches have failed to generate (Chakshusmathi et al., 2004; Varadarajan, Nagarajaram, & Ramakrishnan, 1996). A better understanding of the mechanisms behind the TS phenotype in existing mutants would increase our ability to generate TS mutants via prediction and would provide a resource for the study of essential gene function. To this end, a high throughput method for characterizing the global biophysical effects of temperature-sensitive mutations would increase our ability to link genotype to phenotype in existing, uncharacterized mutants and to develop TS mutants for mechanistic studies of essential genes. Mass spectrometry (MS) based proteomics workflows have evolved to elucidate proteome-wide details and help in the advancement of therapeutic interventions (reviewed in (Doll, Gnad, & Mann, 2019; Mann, Kulak, Nagaraj, & Cox, 2013)). Despite the current capability of MS, there still lacks a high-throughput methodology to evaluate how mutations in a single protein could affect PPI networks, leaving a gap in our ability to predict the phenotypic outcomes of genomic mutations. Previous studies that have obtained high-throughput data on disease-causing mutations, for instance, have relied on approaches such as yeast two-hybrid and chaperone engagement studies which are carried out *in vitro* (Sahni et al., 2015). Although these approaches can provide unique molecular insights, they also rely on gene cloning and overexpression which will artificially alter the balance of allele specific (in diploid cells) and/or PPI partner expression, leading to confounding changes throughout the proteome.

For our model in these experiments, we opted to use TS yeast strains with mutations within individual subunits of the 26S proteasome complex, an essential protein complex responsible for the majority of selective proteolysis occurring in the cell (Coux, Tanaka, & Goldberg, 1996; Hilt & Wolf, 1996). The yeast proteasome is a large multi-subunit protein complex made up of a 20S core complex, consisting of 14 unique subunits, and a 19S regulatory particle, consisting of 19 unique subunits. The proteolytic activity of the proteasome is housed within three subunits of the 20S core proteasome, which is able to degrade substrates prior to complexing with the 19S regulatory particle (reviewed in (Bard et al., 2018)). The 19S regulatory particle functions in substrate recognition, unfolding, and translocation into the core proteasome. Since the proteasome plays a major role in protein turnover and homeostasis, the large-scale changes that could occur in terms of protein abundance levels and protein post-translational modifications, specifically due to the accumulation of polyubiquitin, make it an intriguing model for these studies. One TS mutant was chosen from each of the subcomplexes of the proteasome: one representing the 19S regulatory particle (*rpn5-ts*) and the other representing the 26S proteasome core particle (*pup2-ts*). We performed global proteomics and transcriptomics experiments on these strains and found changes in the abundance of over 1700 proteins and 500 mRNAs, most of which changed similarly across both strains, likely due to improper function of the proteasome. In order to interrogate the biophysical mechanism leading to temperature sensitivity in these strains, we developed a MS-based method with the capability and specificity to measure unique biophysical changes in the thermal stability of the proteome.

The new method described herein is a novel application of thermal proteome profiling (TPP), a recently emerging high-throughput MS strategy that utilizes protein thermal dynamics to characterize such things as target engagement and post-translational modifications (Huang et al., 2019; Jafari et al., 2014; Martinez Molina et al., 2013; Savitski et al., 2014; Tan et al., 2018). We have applied this powerful MS-based platform to yeast strains containing missense mutations in a single essential gene that lead to temperature sensitivity (Kofoed et al., 2015) and assessed changes in the thermal stability of cellular proteins. Missense mutations could alter protein structure and lead to changes in PPIs within associated complexes, and, in theory, affect the thermal stability of either the individual protein or the entire PPI network. In a quest to achieve a high-throughput technology to link phenotype-causing mutations to PPI networks, we hypothesized that TPP could be applied to study the effects of missense mutations on the proteome in a novel method we refer to as <u>Te</u>mperature sensitive (TS) <u>M</u>utant <u>P</u>roteome <u>P</u>rofiling (TeMPP).

TeMPP is a robust method which can be used simultaneously for quantitative measurement of both protein abundance and changes in protein thermal stability between mutants. Interestingly, while about half of the measured proteome changed in abundance, few proteins showed statistically significant changes in their thermal stability, suggesting that proteasome targets (many of them likely polyubiquitylated) do not have major changes in overall stability that could assist in their degradation by the 26S proteasome. RNA sequencing experiments support that these changes are occurring at the level of the proteome and not the transcriptome. Overall, these results show that TeMPP has the capacity to probe both the impact of missense mutations on global proteome thermal stability as well as PPI changes. Furthermore, intersection analysis of protein abundance datasets with thermal profiling and transcriptome measurements gives unique insights into the mechanisms of both mutant protein dysfunction and cell compensation as a consequence of genetic perturbation. The TeMPP method provides foresight into the various potential high-throughput applications of thermal profiling for studying genetic perturbations in diverse cellular systems thereby laying a foundation for novel characterization of the biophysical mechanisms through which mutations lead to protein disfunction and thereby lead to observed phenotypes.

## RESULTS

### Characterizing the global protein and RNA abundance effects of missense mutants in the 26S proteasome

For this work, yeast strains were obtained from a repository of temperature sensitive yeast strains the Hieter lab developed with the intention of creating a resource for studying essential genes in *Saccharomyces cerevisiae* (*S. cerevisiae)* (Kofoed et al., 2015). Random missense mutations were introduced within an essential gene of interest with the resulting strains then screened to identify mutations that caused a temperature sensitive phenotype (Kofoed et al., 2015). The mutated essential genes are expressed from their native promoter and native chromosomal context, allowing for analysis of missense mutation impact on the proteome without the confounding variable of altered protein expression levels. Additionally, the use of haploid yeast strains allows for the characterization of this approach in a genetic system that is not complicated by the presence of multiple alleles for a given gene of interest. These characteristics make these strains an ideal representative model for temperature sensitivity-inducing missense mutations. This work focuses on two strains with missense mutations within subunits of the 26S proteasome: Pup2 (*pup2-ts)*, a 20S proteasome core subunit, and Rpn5 (*rpn5-ts)*, a 19S proteasome regulatory particle subunit.

Global proteomics and polyA RNA sequencing experiments were performed to gain a broad picture of the abundance changes occurring in these strains. Both mutant and wildtype (WT) cells were grown at permissive temperature (30 °C) prior to protein or RNA extraction. For the global proteomics, cells were subjected to the workflow presented in Supp. Fig. 1 and protein was extracted using 8M urea. RNA was extracted using an established hot acid phenol method (Fox, Gao, Smith-Kinnaman, Liu, & Mosley, 2015) prior to library preparation and sequencing on an Illumina system. A total of 3889 proteins and 5750 transcripts were quantified (Supp. Tables 1 & 2) across all three genotypes. In the *pup2-ts* mutant 1731 proteins changed (Fig. 1A) and 702 mRNA transcripts changed in abundance (Fig. 1B). In the *rpn5-ts* mutant, 1997 proteins (Fig. 1C) and 539 mRNA transcripts (Fig. 1D) changed in abundance. Comparisons between protein and mRNA transcript abundance show a low, but positive, correlation (Supp. Fig. 2). With the proteasome’s role in maintaining proteostasis, it is not expected that changes in mRNA transcript abundance would fully explain the changes seen at the level of the proteome in a proteasome mutant background. As both *rpn5-ts* and *pup2-ts* are within the same protein complex, it was unsurprising that both global proteomics and RNA sequencing presented a large number of similar changes between both mutants. There was a 70% overlap in proteins (Supp. Fig. 3) and a 52% overlap in mRNAs (Supp. Fig. 4) that are changing in the two mutants.

**Figure 1:**
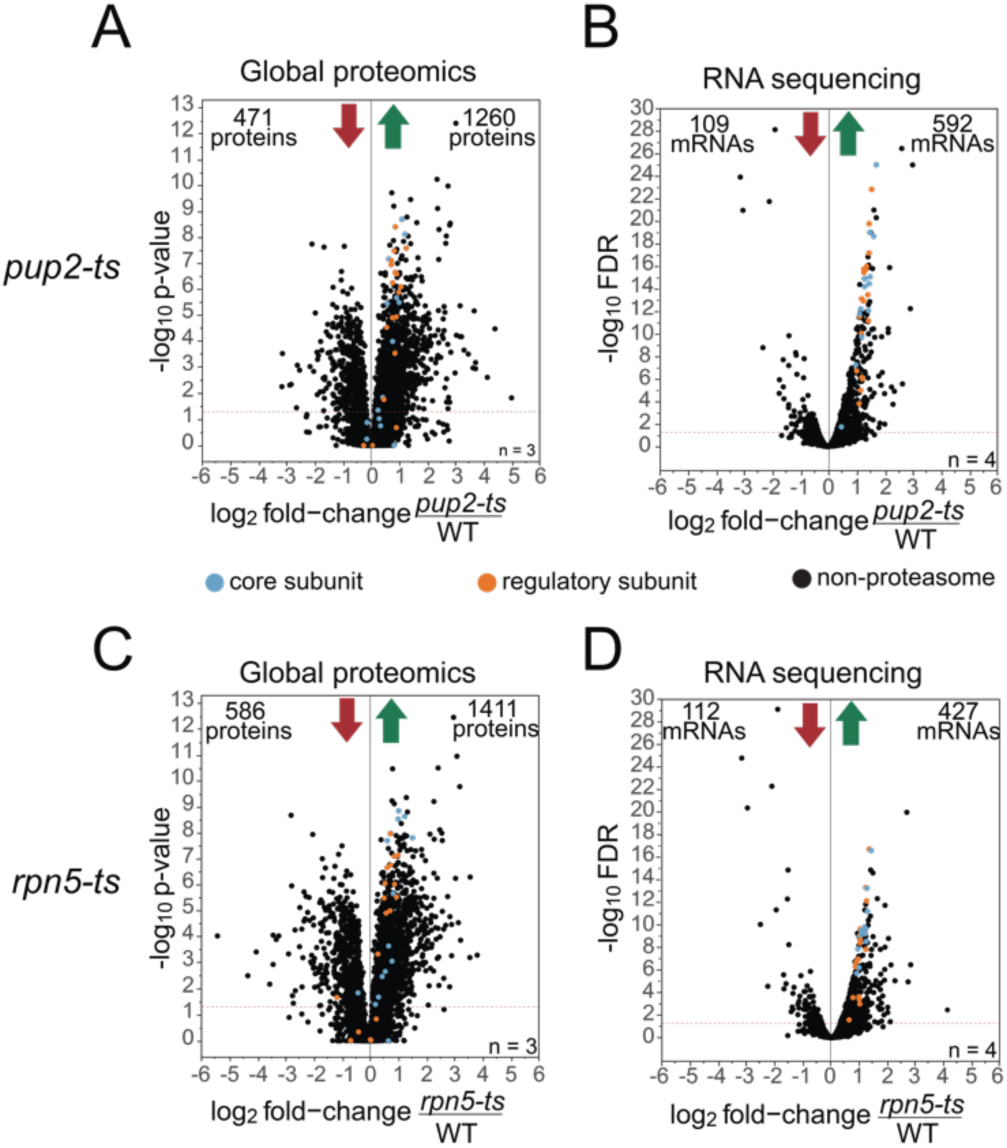
Global changes in protein and mRNA abundance. Volcano plots of fold-change in *pup2-ts/*WT of protein (A) and mRNA transcripts (B) and in *rpn5-ts/*WT of protein (C) and mRNA transcripts (D). X-axis is log_2_ fold-change of mutant/WT; y-axis is -log_10_ *p*-value or FDR. Significance threshold was set at *p*-value or FDR ≤ .05. 3862 data points are shown on each plot. Proteasome core subunits are indicated in blue and regulatory subunits are indicated in orange. A) A total of 1731 proteins significantly changed in *pup2-ts* with 471 proteins decreasing and 1260 proteins increasing in abundance. B) A total of 701 mRNA transcripts significantly changed in *pup2-ts* with 109 transcripts decreasing and 592 transcripts increasing in abundance. C) A total of 1997 proteins significantly changed in *rpn5-ts* with 586 proteins decreasing and 1411 proteins increasing in abundance. D) A total of 539 mRNA transcripts significantly changed in *rpn5-ts* with 112 transcripts decreasing and 427 transcripts increasing in abundance.

Both *pup2-ts* and *rpn5-ts* have increased abundance of many of the proteasome subunits (Supp. Fig. 5) as well as the transcription factor Rpn4 (Supp. Tables 1 & 2). The upregulation at both the mRNA and protein level of the 26S proteasome is a known feedback mechanism for proteasome function that occurs through the transcription factor Rpn4. Rpn4 is a known substrate of the 26S proteasome and increases in abundance as a result of defective proteasome activity (Xie & Varshavsky, 2001). The increase in Rpn4 abundance in both of these strains suggests that it is not being efficiently degraded, strongly indicating that the proteasome is not functioning properly. In an attempt to compensate for decreased proteasome activity in these strains, Rpn4 upregulates the expression of the subunits of the proteasome to increase the number of proteasomes in the cell. Another strong indicator of proteasome dysfunction is an increase of the proteasome chaperone Ump1 in both strains. Ump1 functions in assembly of the core proteasome and is degraded by the 20S proteasome prior to its binding the 19S regulatory particle and formation of the full 26S proteasome (Burri et al., 2000; Ramos, Hockendorff, Johnson, Varshavsky, & Dohmen, 1998). The large increase in Ump1 protein accumulation, 12.7-fold increase in protein relative to a 2.8-fold increase in mRNA, in *pup2-ts* may suggest that the TS-mutations in Pup2 cause a proteasome assembly defect which exacerbates Ump1 accumulation. In *rpn5-ts* the increase in Ump1 protein is comparable to the change in mRNA of *UMP1* (2.5-fold increase in protein and 2.2-fold increase in mRNA), suggesting this may occur through a transcriptional response to the lack of proteasome activity.

### Thermal proteome profiling can be adapted to measure changes resulting from genetic perturbation

Mass spectrometry methods are widely applied to measure changes in protein abundance, protein turnover rates, post-translational modifications (PTMs), and PPIs (Smits & Vermeulen, 2016). Thermal proteome profiling (TPP) is an emerging method that is revolutionizing how MS is able to gain insights into the proteome however it has not yet been applied to biological questions regarding changes that occur as a consequence of genetic perturbation. We have developed an adapted protocol of TPP (Figure 2A) (Franken et al., 2015; Jafari et al., 2014) to address the global effects of missense mutations on the proteome which we have named <u>Te</u>mperature sensitive <u>M</u>utant <u>P</u>roteome <u>P</u>rofiling (TeMPP).

**Figure 2:**
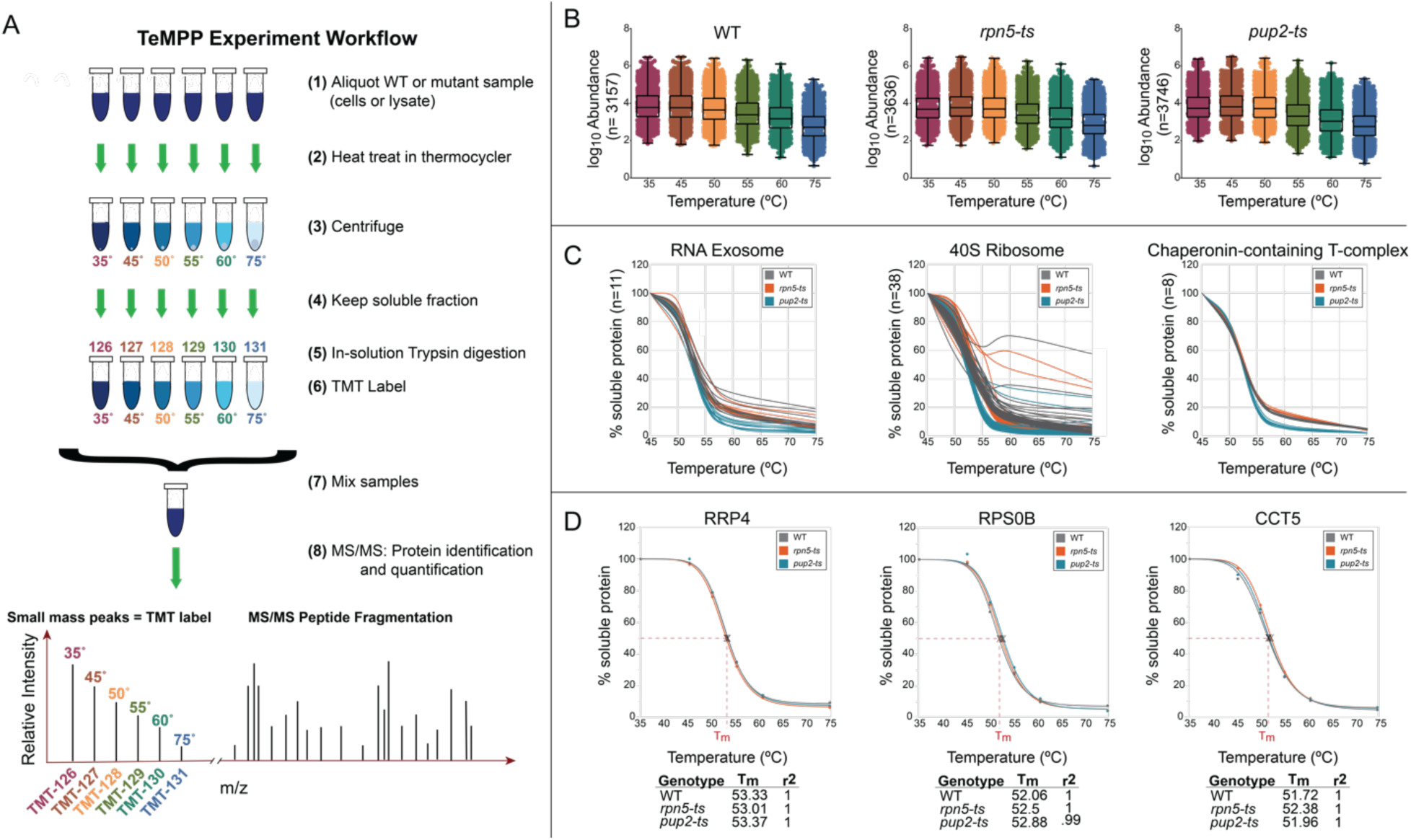
TeMPP can be used to measure changes in PPIs in mutant vs. WT. A) TeMPP workflow. Equal amounts of protein from each lysate was subjected to six different temperature treatments: 35.0 °C; 45.3 °C; 50.1 °C; 55.2 °C; 60.7 °C; and 74.9 °C, to induce protein denaturation. The soluble fractions from each treatment were digested in-solution with Trypsin/Lys-C. Resulting peptides were labeled with isobaric mass tags (TMTsixplex™) and mixed by genotype prior to mass spectrometry (MS) analysis. Resulting MS/MS data were analyzed using Proteome Discoverer™ 2.2 to identify and quantify abundance levels of peptides for each temperature treatment and each genotype. B) Dot plots showing the abundance value for every protein detected from the TemPP experiments in WT, *rpn5-ts,* and *pup2-ts*. Box-and-whiskers plots include minimum, maximum, and median. C) Representative melt curves from a single replicate of the percent soluble protein following heat treatment are shown for selected protein complexes. Proteins isolated from wildtype are shown in gray, *rpn5-ts* in orange, and *pup2-ts* in teal. Shown here are curves of the RNA exosome (11 individual proteins), the 40S ribosome (39 individual proteins), and the Chaperonin-containing T-complex (8 individual proteins). D) Individual protein melt curves were created for every quantified protein and normalized using the TPP R package. T_m_ was calculated as the temperature at which 50% of the protein was denatured. Shown are one protein from each of the complexes in B.

Strains were grown as previously described and lysed in a mild lysis buffer containing 0.1% Tween. Cell lysate from each genotype was subjected to the workflow described in Figure 2A for thermal profiling. Three biological replicates were prepared and were subjected to high pH basic fractionation, resulting in seventy-two LC-MS/MS runs for each genotype. We detected and quantified a total of 4,073 proteins across three biological replicates for each of the three genotypes (Fig. 2B; Supp. Table 3). Data obtained from quantitative MS/MS experiments were used to make melt curves in two different ways (Figures 2C & D). Raw abundances from each temperature treatment were normalized to the ion abundance detected in the 45 °C sample and were plotted in Excel to visualize melt curves of protein complexes in mutant and WT (Figure 2C). Melt curves of three different protein complexes, the RNA exosome, the 40S ribosome, and the Chaperonin-containing T-complex, are shown in Figure 2C. From these we can see that proteins within a complex melt in a similar fashion, consistent with what has been previously shown (Tan et al., 2018). The TPP R package (Childs D, 2018) was used to generate normalized protein melt curves and calculate melt temperatures (T_m_) for ∼3,400-3,600 proteins, depending on the replicate (numerical data for these in Supp. Table 4). Average T_m_ value distributions were plotted for the proteomes of WT as compared to individual mutants (Supp. Fig. 6). Observation of average T_m_ in the range of 50 - 56 °C indicated that the vast majority of proteins denature in this region, inferring the TeMPP strategy using only six temperature points provides appreciably comparable data to that which would have been generated if ten temperature points had been used (Becher et al., 2018; Reinhard et al., 2015; Savitski et al., 2014) and to other methods measuring T_m_ of the proteome (Leuenberger et al., 2017). For the majority of proteins, such as those shown in Figure 2D, we saw very similar T_m_ values between mutant and WT and tight fitting to a sigmoidal curve (as indicated by the r^2^ value). These observations infer a successful adaptation of the TPP workflow for TeMPP with the cost-effective change of using only six temperature points which is of particular advantage for high-throughput screening studies. All biological replicates plots for the proteins shown in Figure 2D are provided in Supp. Fig. 7.

### TeMPP measures the impact of single protein mutants on the stability of the global proteome and on protein-protein interactions

Change in T_m_ (ΔT_m_) values were calculated by taking WT_Tm_ -mutant_Tm_, thereby limiting calculations to proteins detected in both WT and mutant (Supp. Table 5). Further parsing was accomplished by limiting our data to melt curves with r^2^ values > 0.9 and then by proteins that were detected in at least two of the three replicates, providing us with a final analysis of ΔT_m_ in ∼2,050 proteins (Supp. Table 6).

The 19S regulatory particle mutant, *rpn5-ts,* contains 13 missense mutations within Rpn5, which confer temperature sensitivity at 37 °C. Analysis of thermal stability throughout the proteome found significant decreases in the thermal stability of 70 proteins and increases in the stability of 40 proteins (Figure 3A, Supp. Table 7). Analysis of the melt curves of the subunits of the proteasome in *rpn5-ts* showed no changes in the 20S core relative to WT (Figure 3B). Despite the numerous missense mutations within *RPN5*, neither the thermal stability of Rpn5 itself nor any subunits of the 19S regulatory particle were significantly affected (Figure 3C). All replicate data for the raw melt curves for each proteasome subunits are provided in Supp. Fig. 8. Following normalization, thermal melt profiles for the proteasome subunits showed tight reproducibility across all replicates and genotypes indicated that no changes °Ccur in proteasome protein subunit stability in *rpn5-ts* (Figure 3D & E; replicate melt curves of normalized for all proteasome subunits are provided in Supp. Fig. 9). A common mechanism that could explain temperature sensitivity phenotype is that elevated temperature disrupts the structural integrity of a mutant protein. In fact, this is a general concept that is thought to explain temperature sensitivity in many conditional mutant organisms (Lovato et al., 2009). However, these results suggest that the causality of the thermal sensitivity of this strain is not necessarily due to the denaturation of Rpn5 at restrictive temperatures, but likely because of other PPI effects, changes in Rpn5 function, or compensation pathways that adjust for mutant protein function at the permissive temperatures.

**Figure 3:**
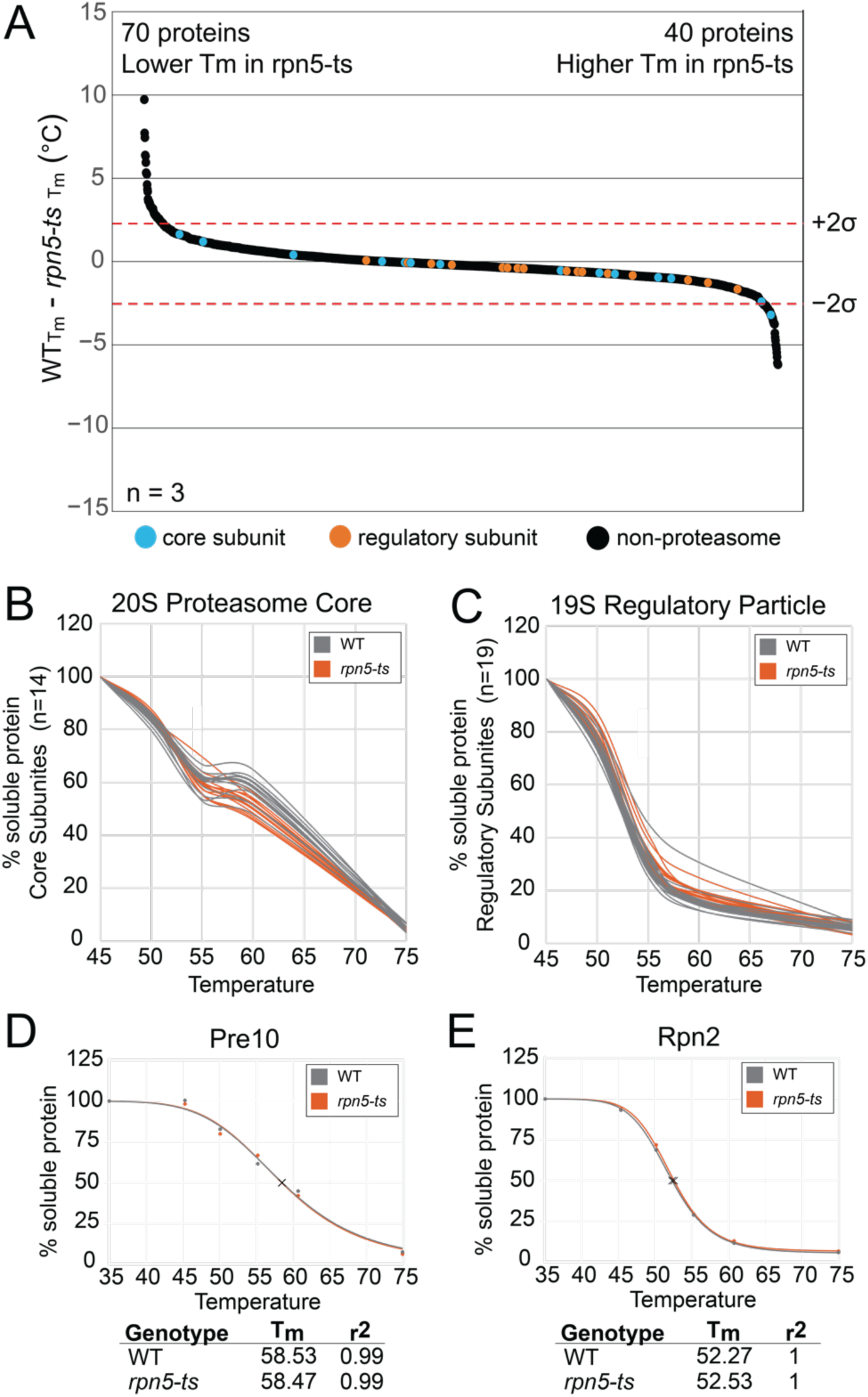
Mutations in Rpn5 do not affect the thermal stability of the proteasome. A) Waterfall plots visualizing whole proteome changes in melt temperature (T_m_), WT-*rpn5-ts*. A total of 2,068 proteins were ordered according to change in T_m_ and plotted. Shown is median values across three biological replicates for proteins that were quantified in at least two replicates. Dotted lines signify a confidence interval of 95%. There were significant decreases in thermal stability of 70 proteins; 40 proteins had significant increases in thermal stability. B & C) Representative melt curves from WT vs. *rpn5-ts* for each of the 14 subunits of the 20S proteasome core (B) and the 19 subunits of the 19S regulatory particle (C) derived from data from one TeMPP experiment. Each line represents an individual subunit. D & E) Individual normalized melt curves of representative protein D) Pre10, from the 20S core and E) Rpn2, from the 19S regulatory particle.

The 20S core particle mutant, *pup2-ts*, contains four missense mutations within Pup2 and is sensitive to growth at 37 °C. A total of 22 proteins were thermally destabilized in the *pup2-ts* mutant, remarkably including all 14 proteasome core subunits (Figure 4; Supp. Table 7). Analysis of the melt curves of each proteasome 19S and 20S subunits showed that in the *pup2-ts* mutant all 14 subunits of the proteasome core are significantly destabilized (Figure 5A). None of the regulatory particle subunits (Figure 5B) had altered thermal stability in *pup2-ts*, suggesting that the mutations in this protein likely affect proteasome activity by disrupting the rigid structure of the core proteasome, and not the more flexible regulatory particle. Normalization of the data via the TPP package resulted in similar findings (Figure 5C & D). The specificity of these findings is quite striking, with nearly 64% of the destabilized proteins °Ccurring within the same protein complex. The normalized data for all replicate melt curves for each proteasome subunit in *pup2-ts* is provided in Supp. Fig. 9. Gene ontology analysis (Mi et al., 2017) of the destabilized protein group identified the GO term proteasomal ubiquitin-independent protein catabolic process as the most enriched with a fold-change enrichment value >100 and a false discovery rate of 4.25e^-25^. These findings clearly show that TeMPP has a very high degree of selectivity for thermally altered proteins and, in this case, their close PPI partners. Interestingly, 103 proteins were significantly stabilized in *pup2-ts* (Figure 4). The *pup2-ts* stabilized proteins could be either directly impacted by the *pup2-ts* mutation or due to secondary impacts of the mutations, such as disruption of proteasome function leading to an accumulation of polyubiquitylation on these proteins or changes in their PPI network.

**Figure 4:**
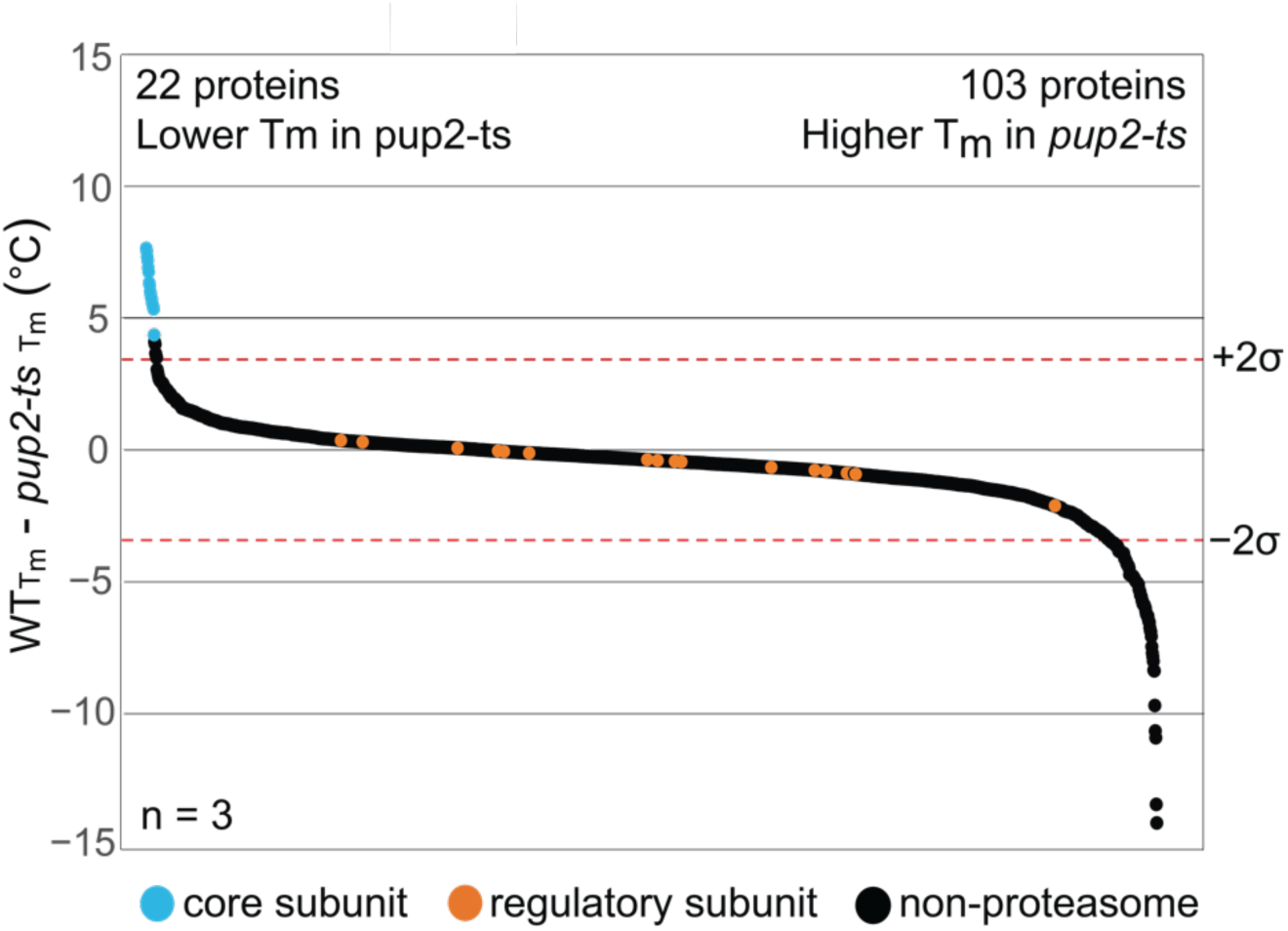
Mutations in Pup2 disrupt the core proteasome. Waterfall plots visualizing whole proteome changes in melt temperature (T_m_), WT-*pup2-ts*. A total of 2,046 proteins were ordered according to change in T_m_ and plotted. Shown is median values across three biological replicates for proteins that were quantified in at least two replicates. Dotted lines signify a confidence interval of 95%. There were significant decreases in thermal stability of 22 proteins; 103 proteins had significant increases in thermal stability.

**Figure 5:**
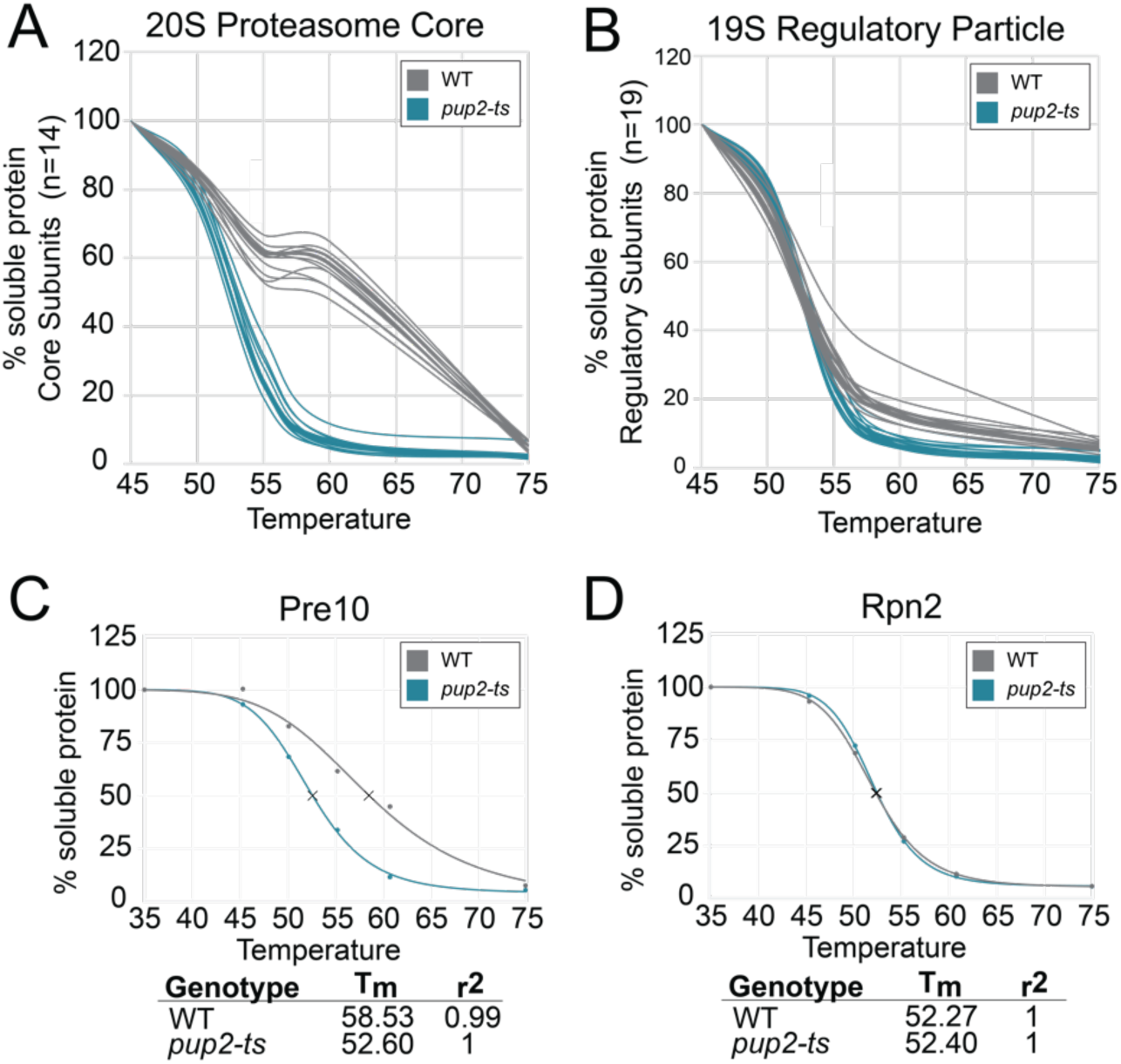
TeMPP uncovers thermal destabilization of all 20S core subunits of the proteasome in *pup2-ts* mutant cells. A & B) Representative raw data melt curves from WT vs. *pup2-ts* for each of the 14 subunits of the 20S proteasome core (A) and the 19 subunits of the 19S regulatory particle (B) derived from data from TeMPP experiment. Each line represents an individual subunit. D & E) Individual normalized melt curves of representative proteins D) Pre10, from the 20S core and E) Rpn2, from the 19S regulatory particle.

### TeMPP can be used to monitor global protein abundance changes in addition to thermal stability

It would be more time effective and less expensive if TeMPP could measure both T_m_s and protein abundance, however previous studies have not directly assessed the reproducibility of the abundances obtained in a TPP-like workflow and their correlation with standard global proteome abundance workflows, which often use denaturing extraction conditions, making them incompatible with TPP. To determine how reproducible protein abundance measurements are from TeMPP, data was compared from the global proteome abundance method and TeMPP. There was a ∼20% difference in the extent of proteome coverage relative to TeMPP, as expected due to difference in extraction buffers. Pearson correlation analysis was performed to compare the measured abundances from the lowest temperature treated sample in the TeMPP experiments to the global TMT proteomics analyses. Despite being isolated in two different buffers, correlation values were greater than 0.8 between the two experiments (Figure 6A) and there was an 82% overlap in protein identification (Figure 6B). This suggests that abundance values obtained from the lowest heat treated TeMPP samples could also be used to measure protein abundance changes with similar depth of proteome coverage, eliminating the need for a separate global proteome abundance experiment and thereby reducing starting material, time, and resources.

**Figure 6:**
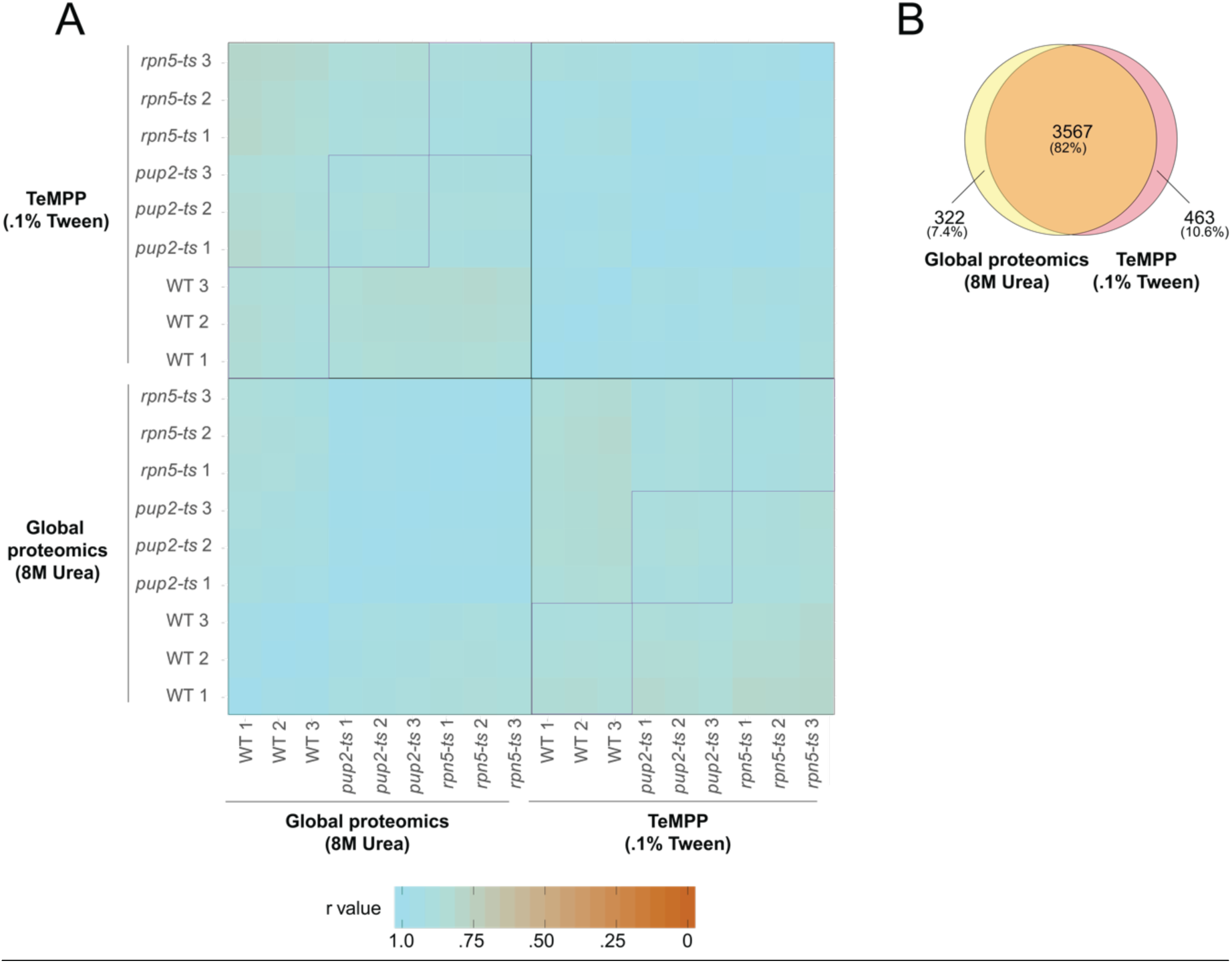
TeMPP can be used to measure global protein abundance. A) Comparison of data acquired from global proteomics experiments (lysis buffer of 8M Urea) vs. TeMPP experiments (lysis buffer containing .1% Tween). Heat map of Pearson correlation values comparing replicates from global proteomics experiments with the lowest temperature of the TeMPP experiments. All correlation values are above 0.80. Indicated sections (purple) highlight comparison of the same genotype across the two methods. B) Venn diagram comparing proteins detected in global proteomics experiments with the TemPP experiments. There is an 82% overlap in quantified proteins between the two methods.

### Multiomics intersection analysis of TeMPP with global proteome abundance and transcriptome data provides unique mechanistic insights into mutant dysfunction

Integration of TeMPP with proteomic and transcriptomic data can provide a clearer picture into the mechanisms linking genotype to phenotype and aid in mechanistic hypothesis generation. Multiomics intersection analysis was used to visualize the total overlap between the TeMPP, RNA sequencing, and global protein abundance data sets through the generation of Upset plots using the UpSetR package in R (Conway, Lex, & Gehlenborg, 2017) and manual dataset intersection analysis (Figure 7). There was little overlap between gene products that are seen to increase or decrease in the same way, and in some cases, there were some that changed in opposite ways, providing potential interesting candidates for further mechanistic studies.

Multiomics intersection analysis of the three datasets revealed subcategories of interest (Figure 7A). For example, in the *pup2-ts* mutant, fourteen of the fifteen proteins seen to be both thermally destabilized and have increased mRNA, intersects of which are indicated in Figure 7A by arrows, are the subunits of the proteasome core, showing the strength of using TeMPP in multiomics intersection analysis for illuminating interesting candidates for mechanism. Aside from the core proteasome subunits, the additional member within this subset is the DNA-damage inducible protein 1 (Ddi1), a ubiquitin binding protein that functions as a proteasome shuttle (reviewed in (Saeki, 2017)). Ddi1 is destabilized in *rpn5-ts* as well, and has been shown to interact with the proteasome via affinity-purification mass spectrometry (Guerrero, Milenkovic, Przulj, Kaiser, & Huang, 2008). It could be that the thermal stability of Ddi1 is dependent on the interaction of Ddi1 with the proteasome or its substrates, and the disruption in proteasome activity is impacting these interactions, leading to Ddi1 destabilization.

In *rpn5-*ts, one particular intersection that stood out in *rpn5-ts* is a gene that was thermally destabilized and decreased in abundance but had an increase in mRNA abundance, Tma17, highlighted by an arrow in Figure 7B. This little-studied protein has been shown to be an important proteasome assembly chaperone in response to stress (Hanssum et al., 2014), and an increase in mRNA suggests the cell maybe trying to compensate for the destabilization and decreased abundance of this protein by upregulating the mRNA transcript. Analyzing the overlap of changes °Ccurring in *pup2-ts* and *rpn5-ts* has the potential for discovery of general proteasome interactors as well as highlighting subunit specific interactors. For example, analyzing the intersect in proteins that are increasing in abundance but whose mRNA levels are not increased in both mutants could be a strategy for the identification of candidate proteasome substrates. A total of 1117 proteins had significant increases in abundance in global proteomics analyses in both *pup2-ts* and *rpn5-ts* (Fig. 8A). In fact, almost 75% of proteins which were found to increase in abundance using global proteomics were found to increase in both mutants (Fig. 8B, abundance increased). A total of 363 mRNA transcripts were upregulated in both *pup2-ts* and *rpn5-ts*, however this was only ∼50% of the total increases observed in mRNA levels in both mutants (Fig. 8). Strikingly, there were a low percentage of proteins that had changes in thermal stability in the same direction in both mutants, 6 destabilized and 3 stabilized (Fig. 8), underscoring the remarkable selectivity and reproducibility of the TeMPP approach. Compared to the commonly applied approaches of transcriptome and proteome abundance analysis, measuring the thermal stability of proteins using TeMPP provided the most unique insights into the underlying impact of each proteasome subunit mutant on the biophysical state of the proteasome and the mechanism of missense mutation dysfunction.

**Figure 7:**
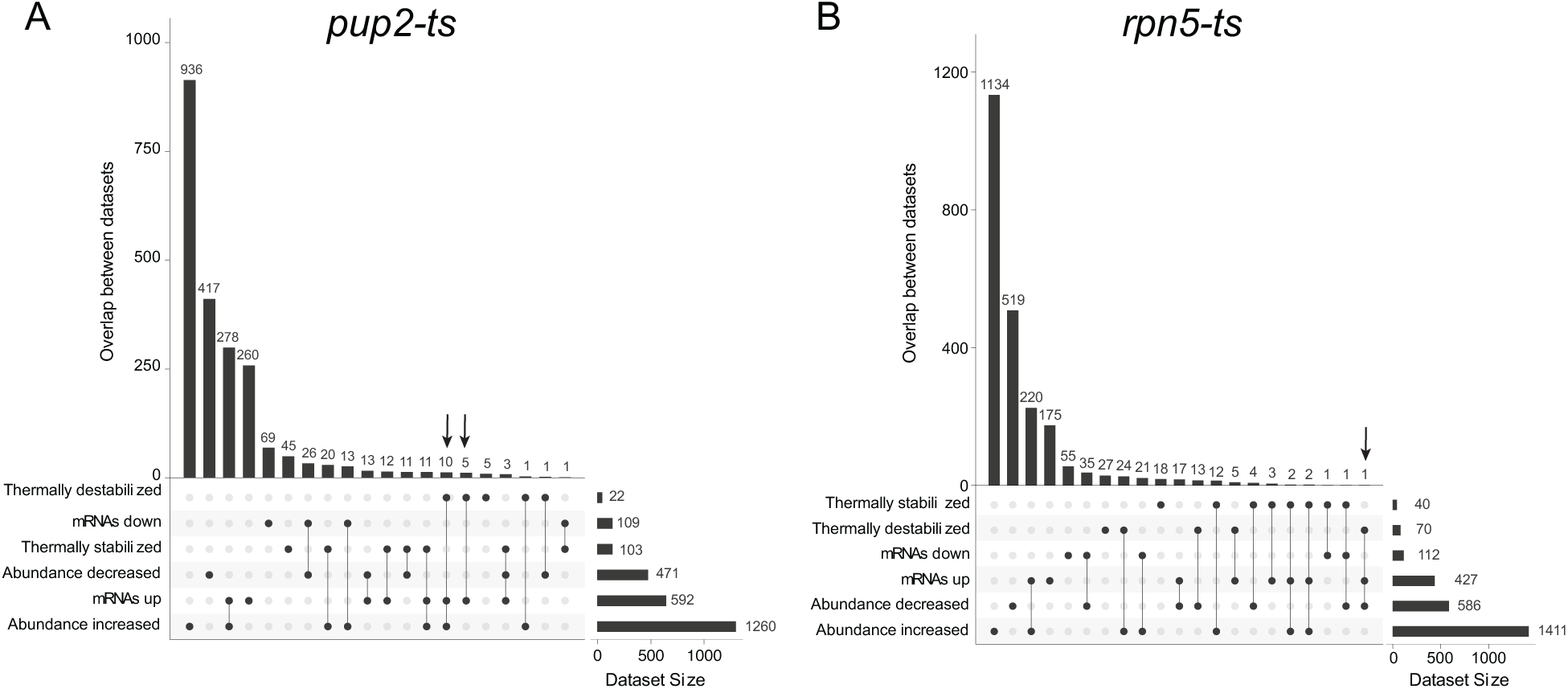
Multiomics intersection analysis of *pup2-ts* and *rpn5-ts*. Upset plots visualizing the overlap of gene products within the sets of significant changes measured in TeMPP, global proteomics, and mRNA sequencing in A) *pup2-ts* and B) *rpn5-ts*. Arrow indicates subset mentioned in the text.

**Figure 8:**
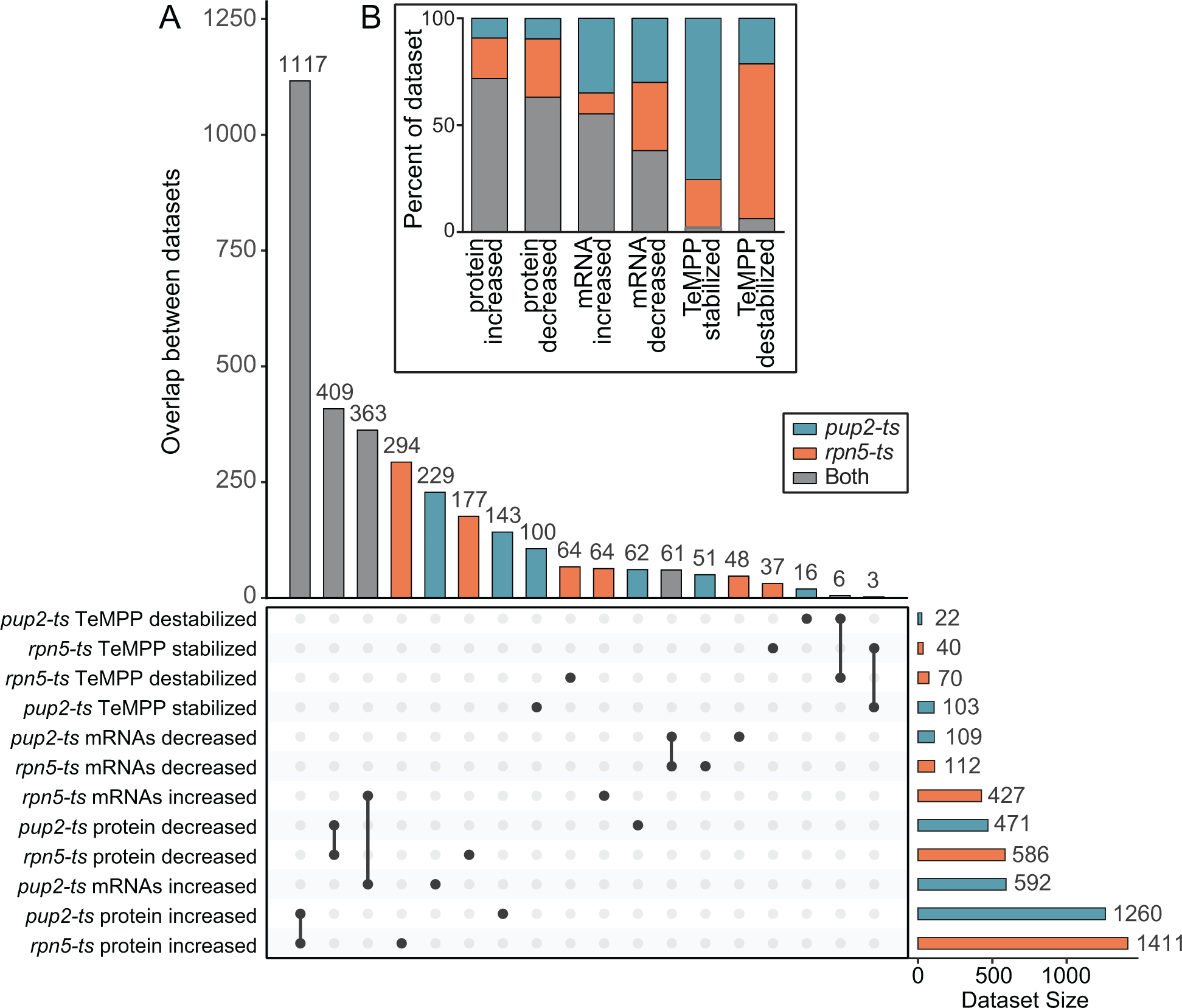
Comparing changes between *rpn5-ts* and *pup2-ts.* Upset plot of the overlap in changes between *rpn5-ts* (orange) and *pup2-ts (*teal). Gene products that are changing in the same way in both genotypes are indicated in gray. B) Stacked bar plot shows the percentage of each data set that is changing in the same direction in both mutants (gray) or changes unique to *rpn5-ts* (orange) or *pup2-ts* (teal).

## DISCUSSION

Global protein abundance measurements and RNA sequencing data in *rpn5-ts* and *pup2-ts* resulted in a large number of possible candidates that could be causing phenotypic changes observed in the mutant strains. Many of the alterations in global protein and RNA abundance were the same across the two mutants, which could lead to the conclusion that they have similar functional disruption. However, the additional data gained through TeMPP on protein stability and potential PPI changes suggest that the *rpn5-ts* and *pup2-ts* mutants are altered through different mechanisms. These missense mutants contain few proteins which are changed in thermal stability, most of which are unique to the mutant subunit studied, suggesting TeMPP to be sensitive enough to measure mutation dependent shifts in thermal stability and protein-specific interactions, even between two proteins within the same protein complex. Use of this method allows for the determination of mechanistic differences in the phenotypes of two mutant strains that would be difficult, or perhaps impossible, to discover through other existing experimental methods in a high-throughput manner. Our study produced quantitative melt curves for over 3,000 proteins across each of the three genotypes evaluated. Though these mutations affect overall proteostasis, only ∼100-130 proteins had a change in thermal stability and there was not a global effect on protein thermal stability, with the average global melt temperature remaining similar to WT. Furthermore, TeMPP identified possible PPIs that are unique to Pup2 and Rpn5 individually, as seen through proteins that change in thermal stability in only one of the two mutants, providing an avenue through which one can study the unique biophysical mechanisms that are °Ccurring due to mutations in different proteins that lead to similar phenotypes.

Performing TeMPP in addition to complementary proteomic and transcriptomic experiments allows for multiomic intersection analysis that may reveal interesting regulatory categories to pursue in follow-up mechanistic interrogations. It remains a possibility that a number of the proteins that increase in abundance in both mutants as a consequence of proteasome perturbation are direct substrates of the proteasome. Additional studies measuring changes in ubiquitylation in these mutant strains may help determine whether some of the changes in protein expression and/or stability in either of the proteasome mutants are due to accumulation of ubiquitylated proteins. Another possibility for an increase in stability in proteins in these mutants is through the actions of cellular chaperone proteins. There was an increase in the abundance of Ssa1 and Ssa2, two Hsp70 family members in both mutants: ∼1.5x increase in *pup2-ts* and a ∼2x increase in *rpn5-ts.* Ssa1/2 have been shown to play a role in delivering misfolded protein to the proteasome during ER-associated degradation (ERAD) (Nakatsukasa, Huyer, Michaelis, & Brodsky, 2008) Additionally, Hsp104, another chaperone linked to ERAD, was seen to increase in protein abundance 1.7x and 1.4x in *pup2-ts* and *rpn5-ts* respectively (Doonan et al., 2019). The up-regulation of cellular chaperones is a potential compensation mechanism for defects in 26S proteasome function.

Our study herein has shown that TeMPP is capable of measuring changes in the thermal stability of proteomes as a result of temperature sensitivity-inducing missense mutations without the need for any additional genetic manipulation or antibody-based interrogation. Considering the inherent challenges with antibody-based methods because of specificity concerns (Baker, 2015), methods that avoid dependence on antibodies, such as TeMPP, are likely to increase the rigor and reproducibility of mutant protein interrogation. Additionally, the lack of a requirement of protein subunit epitope tagging, a genetic manipulation, allows for rapid interrogation of proteome-transcriptome-phenotype characterization. Epitope tagging is a key molecular biology tool for affinity purification followed by mass spectrometry (AP-MS). Although AP-MS is a powerful approach for protein-protein interaction (PPI) identification, it requires a large amount of starting material which is challenging to obtain from many biological systems including patient samples (Dronamraju et al., 2018; Mosley, Florens, Wen, & Washburn, 2009; Mosley et al., 2011). Finally, these data show that TeMPP can be used to reproducibly measure global protein abundance from the low temperature treatment conditions, removing the need for separate global proteomics experiments. This method has the potential to be used across different organisms, even those that are difficult to get large amounts of protein from, to measure systems-level perturbations due to genetic variations.

Not only are mutant proteins important molecular biology tools, but more than 100,000 genomic mutations have been linked to human diseases, of which, ∼60% are predicted to affect protein stability and/or PPIs (Sahni et al., 2015). While genomic and transcriptomic sequencing data are abundant and readily accessible, changes in mRNA have been shown to explain only ∼40% of protein level changes (Schwanhausser et al., 2011). A full understanding of disease relies not only on genomic data, but also on defining the status of cellular proteins in the context of their abundance, post-translational modification status, PPIs, stability, and turnover rate. In addition to the capability of TeMPP for the characterization of TS mutants for studies of normal biological mechanism, use of this method with disease-associated mutant proteins will help lay the foundation for a bioinformatic resource increasing understanding of disease mechanisms and addressing the pathogenicity of missense mutations.

## MATERIALS AND METHODS

### Yeast Strains and Growth Conditions for Proteomics Experiments

Temperature sensitive yeast strains were obtained from the Hieter lab (Kofoed et al., 2015). Two strains with missense mutations within the 26S proteasome, *rpn5-ts* and *pup2-ts*, were chosen for these experiments. Both mutant and wildtype (WT) cells were inoculated at an *A_600_* = 0.3 and grown to an *A_600_* = 0.8 in yeast extract, peptone, dextrose (YPD) medium at permissive temperature (30 °C). YPD was removed by filtration through a nitrocellulose membrane. Cells were flash frozen with liquid nitrogen and stored at -80°C to be used in subsequent sample preparation steps.

### TeMPP Sample Preparation

Cells for each genotype and biologically distinct replicate (i.e. 3 genotypes and 3 biological replicates, for a total of 9), were removed from nitrocellulose membrane using water, pelleted, and re-constituted in lysis buffer containing 40 mM Hepes-KOH (pH 7.5), 10% glycerol, 250 mM NaCl, .1% Tween 20, and fresh yeast protease inhibitors. Resulting suspensions were lysed using a mini bead beater and a vortex genie. After removing the beads lysates were subjected to centrifugation (14,000 x g for 20 minutes) to pellet out cellular debris and to obtain a clear lysate. Protein concentration values for each resulting lysate were determined using the Bio-Rad Protein Assay (Bio-Rad, Hercules, CA) employing vendor provided protocols. All lysates were then diluted to a protein concentration of 5 μg/μl for subsequent heat treatment. Six-50 μl aliquots for each genotype and replicate were then distributed across six PCR tubes, and equilibrated at six temperature points—35.0; 45.3; 50.1; 55.2; 60.7; and 74.9 °C for 3 minutes in a thermocycler (Mastercycler Pro, Eppendorf, Hamberg, Germany) system as described elsewhere (Jafari et al., 2014). Following heat treatment, lysates were centrifuged for twenty minutes at 4 °C to pellet out insoluble protein and then to decant the soluble fraction. A 20 μL aliquot from each heat-treated sample was subject to TCA (Trichloroacetic acid) based protein precipitation overnight. Following centrifugation and acetone-based wash steps, resulting protein pellets were reconstituted in 30 μL of 8M urea in Tris-HCl (pH 8.0). Samples were subjected to reduction of Cys-Cys bonds with 5 mM tris (2-carboxyethyl) phosphine hydrochloride (TCEP), and alkylation with 10 mM chloroacetaminde (CAM) to protect the reduced Cys residues. Samples were diluted to 2M urea and were digested in-solution overnight using Trypsin (Promega Corporation, Madison, WI) as previously described to derive peptides (Mosley et al., 2009; Mosley et al., 2011; Smith-Kinnaman et al., 2014). Samples were de-salted using a Sep-Pak® Vac 1cc C18 Cartridge (Waters Corporation, Milford, MA) according to the vendor provided vacuum manifold protocol that involves: 1) priming the column 2) loading the peptides on to column 3) washing the column to remove any buffers from the immobilized peptides 4) elution of immobilized peptides. This was performed to remove buffer additives and excess reagents as many lysis buffers are known to interfere with the TMT labeling reaction. Resulting peptide elutions were dried using a speed vacuum and subjected to TMT labelling using vendor provided protocols. A sixplex kit (Thermo Scientific™, Waltham, MA) was employed for the purpose and, more specifically, channels— TMT126; TMT127; TMT128; TMT129; TMT130 and TMT131 were respectively used to label peptide solutions derived from the 35.0; 45.3; 50.1; 55.2; 60.7; and 74.9 °C temperature treatments. Labeling reactions were quenched and the six labeled samples were mixed and were dried using a speed vacuum system. Dried samples were fractionated using reversed phase fractionation columns (8 fractions) employing vendor provided protocols (Pierce Biotechnology, Waltham, MA). The resulting 8 fractions were dried using a speed vacuum system and resuspended in .1% formic acid (30 μL) prior to nano-LC-MS/MS analysis as described below.

### TeMPP data acquisition – nano-LC-MS/MS

Nano-LC-MS/MS analyses were performed on a Q-Exactive Plus™ mass spectrometer (Thermo Scientific™, Waltham, MA) coupled to an EASY-nLC™ HPLC system (Thermo Scientific™, Waltham, MA). Ten μL equivalent volume of the resuspended fractions from above were loaded using 300 bar as applied maximum pressure onto an in-house prepared reversed phase column. Each reversed phase column was prepared by pulling a 100 μm fused-silica column to carry *ca.* 5 μm tip for the nanospray using a P-2000 laser puller, and then packing the capillary with C18 reverse phase resin (particle size: 3 μm diameter; Dr. Maisch HPLC GmbH, Ammerbuch, Germany). The peptides were eluted using a varying mobile phase (MP) gradient from 95% phase A (FA/H2O 0.1/99.9, v/v) to 24% phase B (FA/ACN 0.4/99.6, v/v) for 150 mins, from 24% phase B to 35% phase B for 25 mins and then keeping the same MP-composition for 5 more minutes at 400 nL/min to ensure elution of all peptides. Nano-LC mobile phase was introduced into the mass spectrometer using a Nanospray Flex Source (Proxeon Biosystems A/S, Denmark). The heated capillary temperature was kept at 275 °C and ion spray voltage was kept at 2.5 kV. During peptide elution, the mass spectrometer method was operated in positive ion mode for 180 minutes, programmed to select the most intense ions from the full MS scan using a top 20 method. Additional parameters: Microscans 1; Resolution 70k; AGC target 3E6; Maximum IT 50 ms; Number of scan ranges 1; Scan range 400 to 1600 m/z; and Spectrum data type “profile”, and then to perform data dependent MS/MS scans with parameters: Microscans 1; Resolution 35k; AGC target 1E5; Maximum IT 64 ms; Loop count 20; MSX count 1; Isolation window 0.7 m/z; Fixed first mass 100 m/z; NCE 38.0; and Spectrum data type “Centroid”. The respective data dependent settings were set with parameters: Minimum AGC target of 1.00e3; Intensity threshold of 1.6e4; Apex trigger as “-”; Charge exclusion as “1,7,8, >8“; Multiple Charge States as “all”; Peptide match as “preferred”; Exclude isotopes as “on”; Dynamic exclusion of 30.0 s; If idle “pick others”. The data were recorded using Thermo Xcalibur (4.1.31.9) software (Copyright 2017, Thermo Fisher Scientific Inc.).

### Sample preparation for global quantitative proteomics comparison of mutant proteomes relative to WT

Three biological replicates of WT, *pup2-ts,* and *rpn5-ts* were prepared as explained above. Unlike the previous preparation, cells were lysed in 8M urea using a mini bead beater and a vortex genie. 50 μg equivalent of protein from each lysate were then subjected to reduction, alkylation, proteolytic digestion, and de-salting as described above. Peptide samples were labeled using a 10plex kit (Thermo Scientific™, Waltham, MA). Specific labelling reagents corresponding to TMT127N; TMT127C; TMT128N; TMT128C; TMT129N; TMT129C; TMT130N, TMT130C, and TMT131 were respectively used to label the peptide solutions derived from three replicates from each genotype *WT*, *PUP2-TS*, and *RPN5-TS*. Following reaction quenching, mixing, and the subsequent drying step, the resulting peptide mixture was fractionated as described, resulting in 8 fractions. Each fraction was dried completely using a speed vacuum system and resuspended in .1% formic acid (30 μL) prior to the below explained nano-LC-MS/MS analysis.

### Global proteomics data acquisition – nano-LC-MS/MS

Nano-LC-MS/MS analyses were performed on an Orbitrap Fusion™ Lumos™ mass spectrometer (Thermo Scientific™, Waltham, MA) coupled to an EASY-nLC™ HPLC system (Thermo Scientific™, Waltham, MA). 18 μL equivalent volume of the resuspended fractions were loaded onto an in-house prepared reversed phase column using 600 bar as applied maximum pressure. Each reversed phase column was prepared by pulling a 100 μm fused-silica column to carry *ca.* 5 μm tip for the nanospray using a P-2000 laser puller, and then packing the capillary with C18 reverse phase resin (particle size: 3 μm diameter; Dr. Maisch HPLC GmbH, Ammerbuch, Germany). The peptides were eluted using a 180-minute gradient increasing from 95% buffer A (0.1% formic acid in water) and 5% buffer B (0.1% formic acid in acetonitrile) to 25% buffer B at a flow rate of 400 nL/min. The peptides were eluted using a 180-minute gradient increasing from 95% buffer A (0.1% formic acid in water) and 5% buffer B (0.1% formic acid in acetonitrile) to 25% buffer B at a flow rate of 400 nL/min. Nano-LC mobile phase was introduced into the mass spectrometer using a Nanospray Flex Source (Proxeon Biosystems A/S, Denmark). During peptide elution, the heated capillary temperature was kept at 275 °C and ion spray voltage was kept at 2.6 kV. The mass spectrometer method was operated in positive ion mode for 180 minutes having a cycle time of 3 seconds for MS/MS acquisition.

MS data was acquired using a data-dependent acquisition method that was programmed to have 2 data dependent scan events following the first survey MS scan. During MS1, using a wide quadrupole isolation, survey scans were obtained with an Orbitrap resolution of 120 k with vendor defined parameters―m/z scan range, 375-1500; maximum injection time, 50; AGC target, 4E5; micro scans, 1; RF Lens (%), 30; “DataType”, profile; Polarity, Positive with no source fragmentation and to include charge states 2 to 7 for fragmentation. Dynamic exclusion for fragmentation was kept at 60 seconds. During MS2, the following vendor defined parameters were assigned to isolate and fragment the selected precursor ions. Isolation mode = Quadrupole; Isolation Offset = Off; Isolation Window = 0.7; Multi-notch Isolation = False; Scan Range Mode = Auto Normal; FirstMass = 120; Activation Type = CID; Collision Energy (%) = 35; Activation Time = 10 ms; Activation Q = 0.25; Multistage Activation = False; Detector Type = IonTrap; Ion Trap Scan Rate = Turbo; Maximum Injection Time = 50 ms; AGC Target = 1E4; Microscans = 1; DataType = Centroid. During MS3, daughter ions selected from neutral losses (e.g. H_2_O or NH_3_) of precursor ion CID during MS2 were subjected to further fragmentation using higher-energy C-trap dissociation (HCD) to obtain TMT reporter ions and peptide specific fragment ions using following vendor defined parameters. Isolation Mode = Quadrupole; Isolation Window =2; Multi-notch Isolation = True; MS2 Isolation Window (m/z) = 2; Number of notches = 3; Collision Energy (%) = 65; Orbitrap Resolution = 50k; Scan Range (m/z) = 100-500; Maximum Injection Time = 105 ms; AGC Target = 1E5; DataType = Centroid. The data were recorded using Thermo Scientific Xcalibur (4.1.31.9) software (Copyright 2017 Thermo Fisher Scientific Inc.).

### Protein identification and quantification

Resulting RAW files were analyzed using Proteome Discover™ 2.2 (Thermo Scientific™, Waltham, MA). The SEQUEST HT search engine was used to search against a yeast protein database from the UniProt sequence database (December 2015). Specific search parameters used were: trypsin as the proteolytic enzyme, peptides with a max of 2 missed cleavages, precursor mass tolerance of 10 ppm, and a fragment mass tolerance of 0.6 Da. Static modifications used for the search were, 1) carbamidomethylation on cysteine(C) residues; 2) TMTsixplex label on lysine (K) residues and the N-termini of peptides. Dynamic modifications used for the search were oxidation of methionine and acetylation of N-termini. Percolator False Discovery Rate was set to a strict setting of 0.01 and a relaxed setting of 0.05. Values from both unique and razor peptides were used for quantification. For the TeMPP experiments, no normalization setting was used for protein quantification; for the global protein abundance measurements peptides were normalized by “total peptide amount”.

### Thermal proteome profiling (TPP) analysis and waterfall plots

The TPP package (v3.12.0) (Childs D, 2018) in R was used to generate normalized melt curves and to determine protein melt temperatures as described previously (Franken et al., 2015). Changes in T_m_ for each protein were calculated utilizing the function WT (T_m_) – mutant (T_m_), where WT (T_m_) represents the median melt temperature of a protein determined in WT samples and mutant (T_m_) represents the median melt temperature of a protein determined in mutant samples. Resulting data processing was performed in R Studio (R Studio for Mac, Version 1.1.456). Data was parsed down to proteins that were detected in at least two of the three replicates and had melt curves with r^2^ values >0.9. Proteins were ranked according to median change in T_m_ and ordered from the largest change (proteins that were destabilized in the mutant) to smallest change (proteins that were stabilized in the mutant). From the distribution of T_m_ changes in a single comparison, T_m_ changes that are out of the range, mean (T_m_ change) ± 2**σ** (**σ** being the standard deviation), were considered statistically significant T_m_ changes, and identified as proteins destabilized or stabilized due to the mutation. Waterfall plots were created using *ggplot2* (Mi et al., 2017).

### RNA extraction

Both mutant and WT cells were inoculated at an *A_600_* = 0.5 in YPD medium and permitted to recover for 90 minutes at permissive temperature (30 °C). Cells were pelleted, washed with water, and resuspended in 10ml Acetate-EDTA (AE) buffer (50mM sodium acetate pH 5.2, 10mM EDTA). RNA was isolated using a hot acid phenol method as described previously (Fox et al., 2015). In brief, 800 μl 20% SDS and 10ml cold acid phenol were added to each sample in a Nalgene phenol-resistant tube. Samples were incubated for ten minutes with rotation in a hybridization oven at 65°C and cooled on ice for 5 minutes. The samples were then centrifuged at 10,000 rpm for 15 minutes and the top phase was transferred to a 50 ml 5 PRIME Phase Lock Gel Tube (Ref #2302870). 13 ml chloroform was added and samples were centrifuged at 3000 rpm for 10 minutes; the remaining top phase was poured into a new phenol-resistant tube. 1/10 volume sodium acetate at pH 5.2 and equal volume isopropanol was added and precipitated RNA was collected by centrifugation at 12,000 rpm for 45 minutes. The RNA pellet was washed with 70% EtOH and dried in a fume hood. RNA was resuspended with molecular biology grade water. Following purification, RNA was treated using the TURBO DNA-free kit (Thermo Scientific™, Waltham, MA) to remove DNA contamination.

### Library Preparation and sequencing

The concentration and quality of total RNA samples was first assessed using Agilent 2100 Bioanalyzer. A RIN (RNA Integrity Number) of five or higher was required to pass the quality control. Then one hundred nanograms of RNA per sample were used to prepare dual-indexed strand-specific cDNA libraries using the KAPA mRNA HyperPrep kit (Roche). The resulting libraries were assessed for quantity and size distribution using Qubit and Agilent 2100 Bioanalyzer. Pooled libraries were utilized for clustering amplification on cBot using HiSeq 3000/4000 PE Cluster Kit and sequenced with 2×75bp paired-end configuration on HiSeq4000 (Illumina) using a HiSeq 3000/4000 PE SBS Kit. A Phred quality score (Q score) was used to measure the quality of sequencing. More than 94% of the sequencing reads reached Q30 (99.9% base call accuracy).

### Sequence alignment and gene counts

The sequencing data were first assessed using FastQC (v.0.11.5, Babraham Bioinformatics, Cambridge, UK) for quality control. All sequenced libraries were mapped to the yeast genome (UCSC sacCer3) using STAR RNA-seq aligner (v.2.5) (Dobin et al., 2013) with the following parameter: “--outSAMmapqUnique 60”. The reads distribution across the genome was assessed using bamutils (from ngsutils v.0.5.9) (Breese & Liu, 2013). Uniquely mapped sequencing reads were assigned to S288C R64-2-1 20150113 annotated genes using featureCounts (subread v.1.5.1) (Liao, Smyth, & Shi, 2014) with the following parameters: “-s 2 –p –Q 10”. Genes with read count per million (CPM) < 0.5 in ≥ 4 of the samples were removed. The data was normalized using TMM (trimmed mean of M values) method. Differential expression analysis was performed using edgeR (v.3.12.1) (McCarthy, Chen, & Smyth, 2012; Robinson, McCarthy, & Smyth, 2010). False discovery rate (FDR) was computed from p-values using the Benjamini-Hochberg procedure.

### Global abundance plots

Volcano plots and correlation plots were created in R Studio using *ggplot2.* Abundance values, fold-change ratios and *p*-values for the proteomics data were calculated by Proteome Discover™ 2.2 (Thermo Scientific™, Waltham, MA). FPKM values, fold-change ratios and FDR values were calculated as described above. *p*-value and FDR cutoff for significance was set at < 0.05. Venn diagram was created using Venny 2.1 (Oliveros, 2007-2015). Abundance dot plots were created using GraphPad Prism version 6.00 for Windows. Upset plots were created using the UpSetR package in R Studio (Conway et al., 2017). ADD IN INFORMATION FOR OVERLAP ANALYSIS STATS HERE.

## ACKNOWLEDGEMENTS

We would like to thank the current members of the Mosley lab: Whitney Smith-Kinnaman, Katlyn Hughes Burriss, and Dominique Baldwin and the IUSM proteomics core: Emma Doud. We would also like to thank Maureen Harrington and Mark Goebl for helpful discussions of this work. A portion of the funding for this project was provided by National Institute of Health T32 HL007910 (SAPJ). Reagents to continue this work have graciously been provided via the Thermo Scientific TMT Research Award. This project was supported, in part, with support from the Indiana Clinical and Translational Sciences Institute funded, in part by Award Number UL1TR002529 from the National Institutes of Health, National Center for Advancing Translational Sciences, Clinical and Translational Sciences Award. Acquisition of the IUSM Proteomics core instrumentation used for this project was provided by the Indiana University Precision Health Initiative.

## AUTHOR CONTRIBUTIONS

S.A.P.J.: obtained funding for the project, designed and performed biochemical experiments, analyzed data (mass spectrometry and intersection analysis), prepared the figures, and wrote the manuscript. G.Q. prepared samples for LC-MS/MS analysis and contributed to protocol development discussions. H.R.S.W. participated in computational analysis of the mass spectrometry experiments and use of R and provided input on writing of the manuscript. S.A.P.J. and J.F.V. performed the RNA sequencing experiments. E.R.S. performed computational analysis of the RNA sequencing data. ABW: contributed to the design of experiments, data analysis, and writing of the manuscript. A.L.M.: Oversaw various aspects of the project and provided funding for the project, provided direction on data analysis and figure preparation, and wrote the manuscript.

## DATA AVAILABILITY

All analyzed datasets used for this study are referenced in the text and included in the supplementary information. Raw datasets from the proteomics experiments are available from the corresponding authors upon reasonable request. The RNAseq data deposition to Gene Expression Omnibus (GEO) is pending.

## CODE AVAILABILITY

Any code used for data analysis is available as referenced. Minor code changes made for application to the data analyzed are available upon request.

---

**Supplemental Figure 1:**
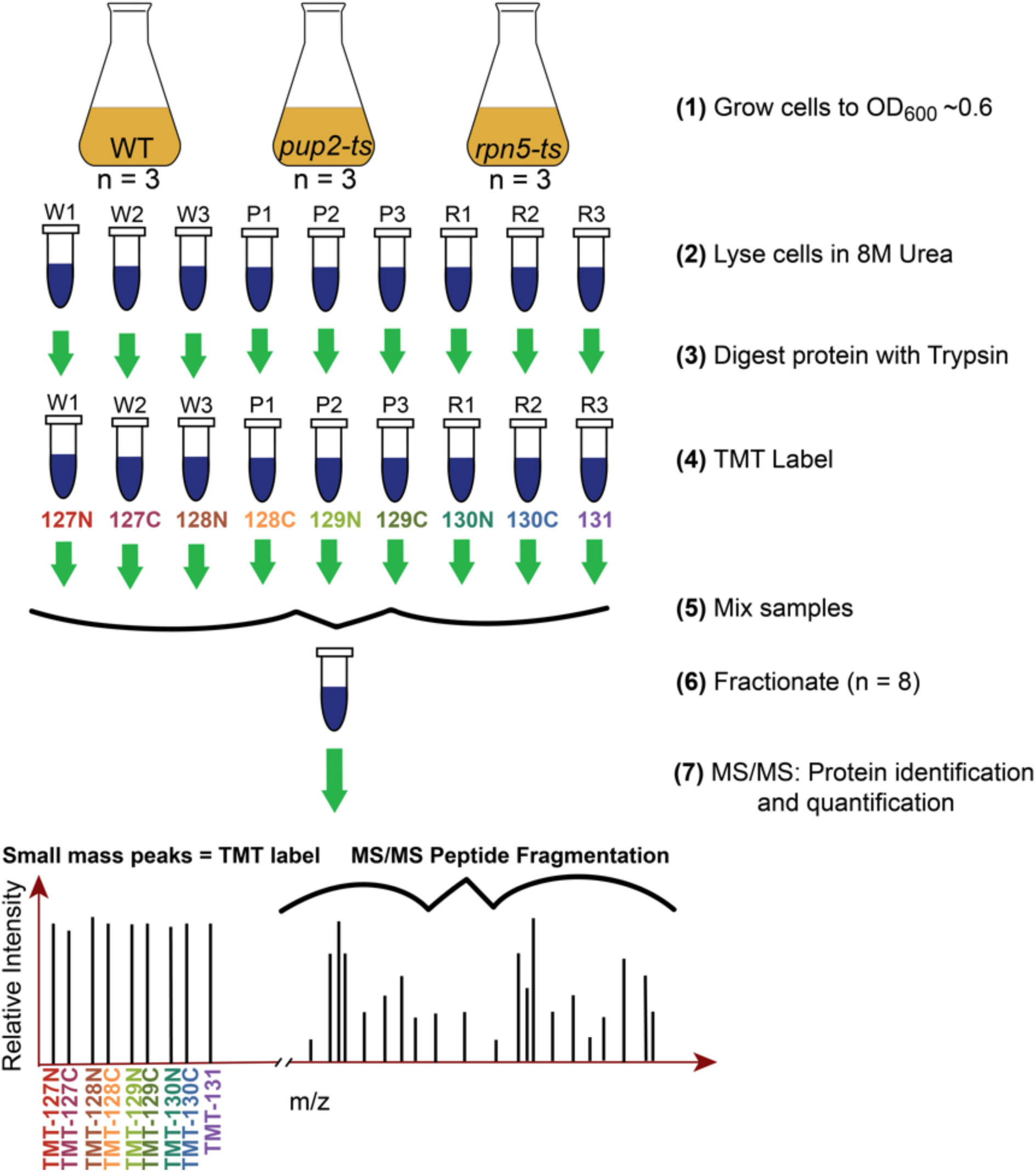
Workflow for quantitative measurement of global protein abundance. Three biological replicates of WT and mutant lysates were subjected to lysis in 8M urea. Equal amounts of protein from each sample were digested with Trypsin/Lys-C and resulting peptides were labeled with isobaric mass tags (TMT10plex™). All nine labeled samples were mixed and fractionated prior to analysis on a Q-Exactive Plus™. Mass spectra were analyzed using Proteome Discoverer™ 2.2 to identify and quantify abundance levels of peptides for each genotype.

**Supplemental Figure 2:**
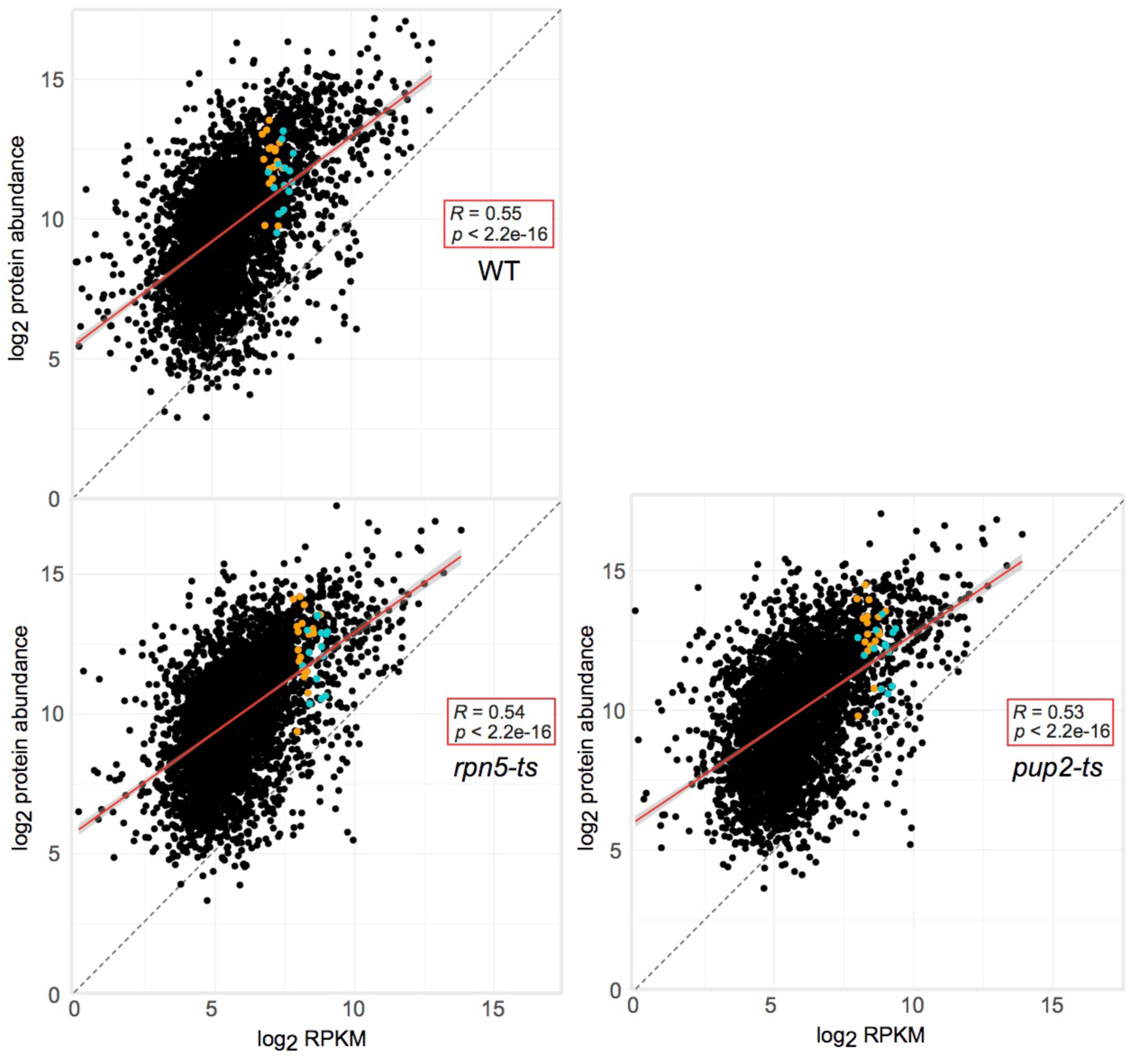
Log-log plots. Log-log plots comparing protein and mRNA abundance in WT, *rpn5-ts,* and *pup2-ts.* All three genotypes show a weak positive correlation with an *R* value ∼0.55. Gray dotted line represents y = x. Red line is the linear regression. Proteasome core subunits are indicated in blue and regulatory subunits are indicated in orange.

**Supplemental Figure 3 (part A/B):**
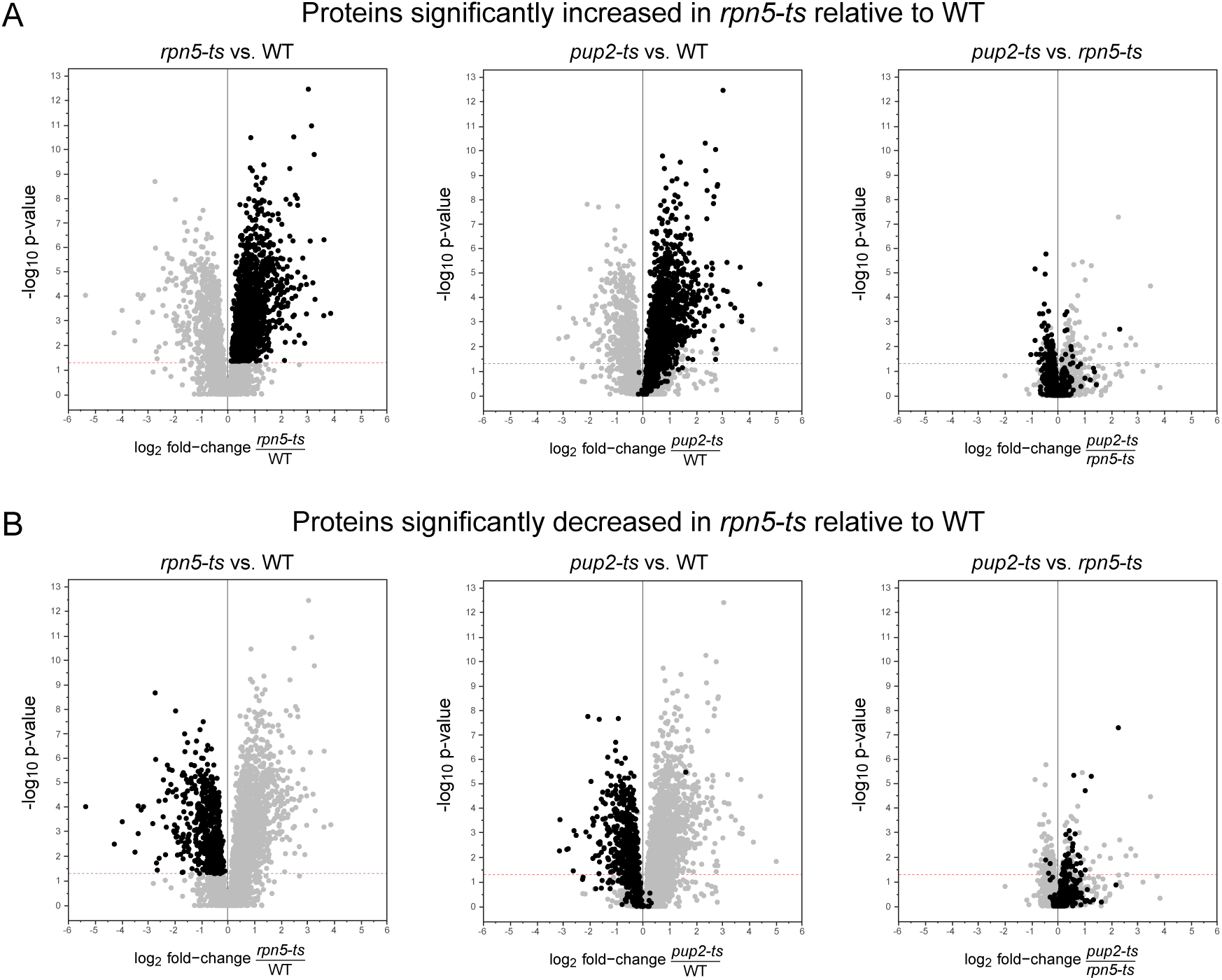
Comparing protein abundance changes in each genotype. Volcano plots of protein quantification from global TMT analysis of WT, *rpn5-ts,* and *pup2-ts*. X-axis is -log_10_ *p*-value; y-axis is log_2_ fold-change of mutant/WT or mutant/mutant (n=3). Significance threshold was set at *p*-value ≤ .05. A total of 3889 proteins are shown on each plot. Each panel shows the three comparative plots shown in the main text but with highlighted A) proteins increased in *rpn5-ts* relative to WT, B) proteins decreased in *rpn5-ts* relative to WT

**Supplemental Figure 3 (part C/D):**
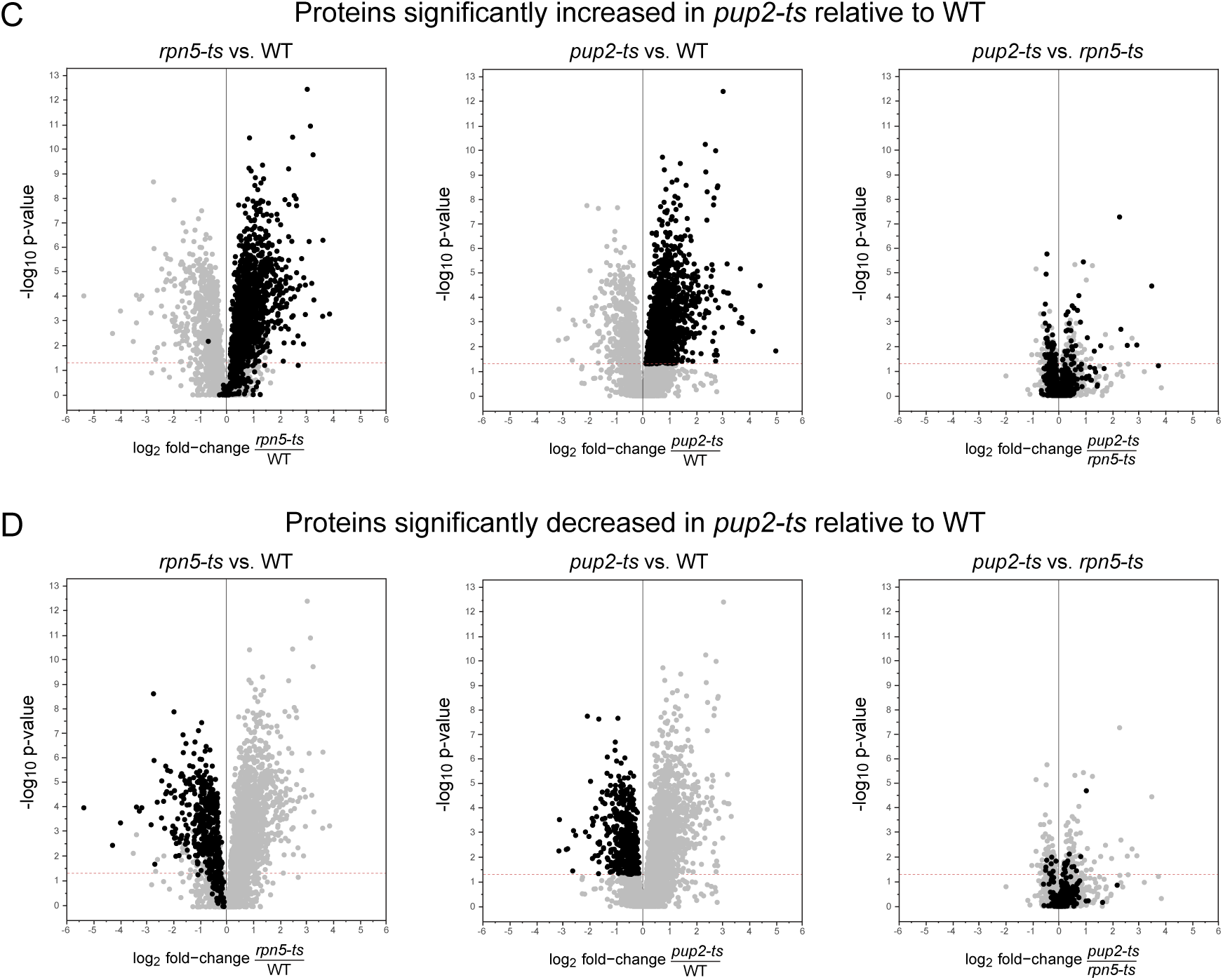
Comparing protein abundance changes in each genotype. Volcano plots of protein quantification from global TMT analysis of WT, *rpn5-ts,* and *pup2-ts*. X-axis is -log_10_ *p*-value; y-axis is log_2_ fold-change of mutant/WT or mutant/mutant (n=3). Significance threshold was set at *p*-value ≤ .05. A total of 3889 proteins are shown on each plot. Each panel shows the three comparative plots shown in the main text but with highlighted C) proteins increased in *pup2-ts* relative to WT, and D) Proteins decreased in *pup2-ts* relative to WT.

**Supplemental Figure 4 (part A/B):**
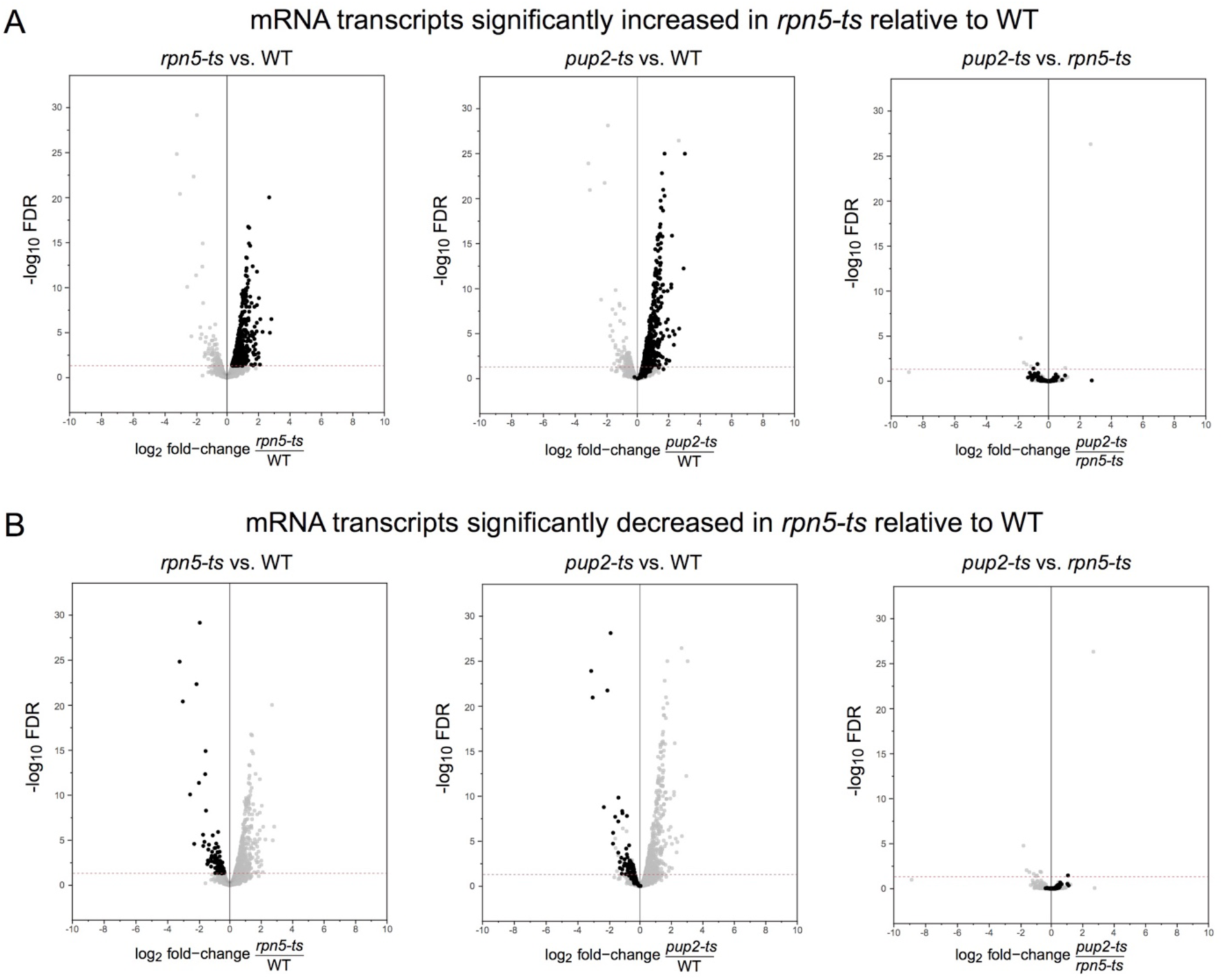
Comparing protein abundance changes in each genotype. Volcano plots of mRNA transcripts global RNAseq analysis of WT, *rpn5-ts,* and *pup2-ts*. X-axis is -log_10_ *FDR*; y-axis is log_2_ fold-change of mutant/WT or mutant/mutant (n=3). Significance threshold was set at FDR ≤ .05. A total of 3862 proteins are shown on each plot. Each panel shows the three comparative plots shown in the main text but with highlighted A) proteins increased in *rpn5-ts* relative to WT, B) proteins decreased in *rpn5-ts* relative to WT

**Supplemental Figure 4 (part C/D):**
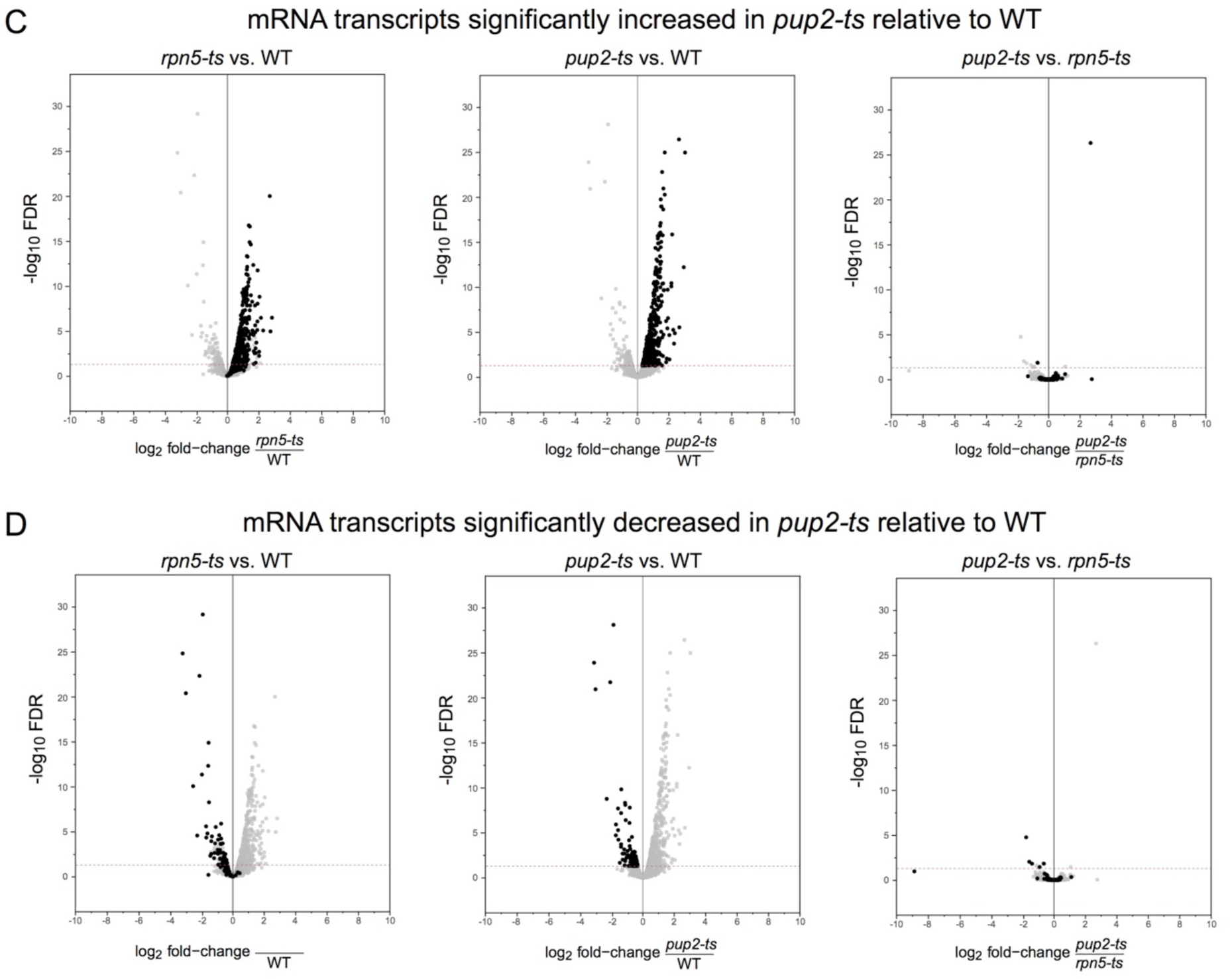
Comparing protein abundance changes in each genotype. Volcano plots of mRNA transcripts global RNAseq analysis of WT, *rpn5-ts,* and *pup2-ts*. X-axis is -log_10_ *FDR*; y-axis is log_2_ fold-change of mutant/WT or mutant/mutant (n=3). Significance threshold was set at FDR ≤ .05. A total of 3862 proteins are shown on each plot. Each panel shows the three comparative plots shown in the main text but with highlighted C) proteins increased in *pup2-ts* relative to WT, and D) Proteins decreased in *pup2-ts* relative to WT.

**Supplemental Figure 5:**
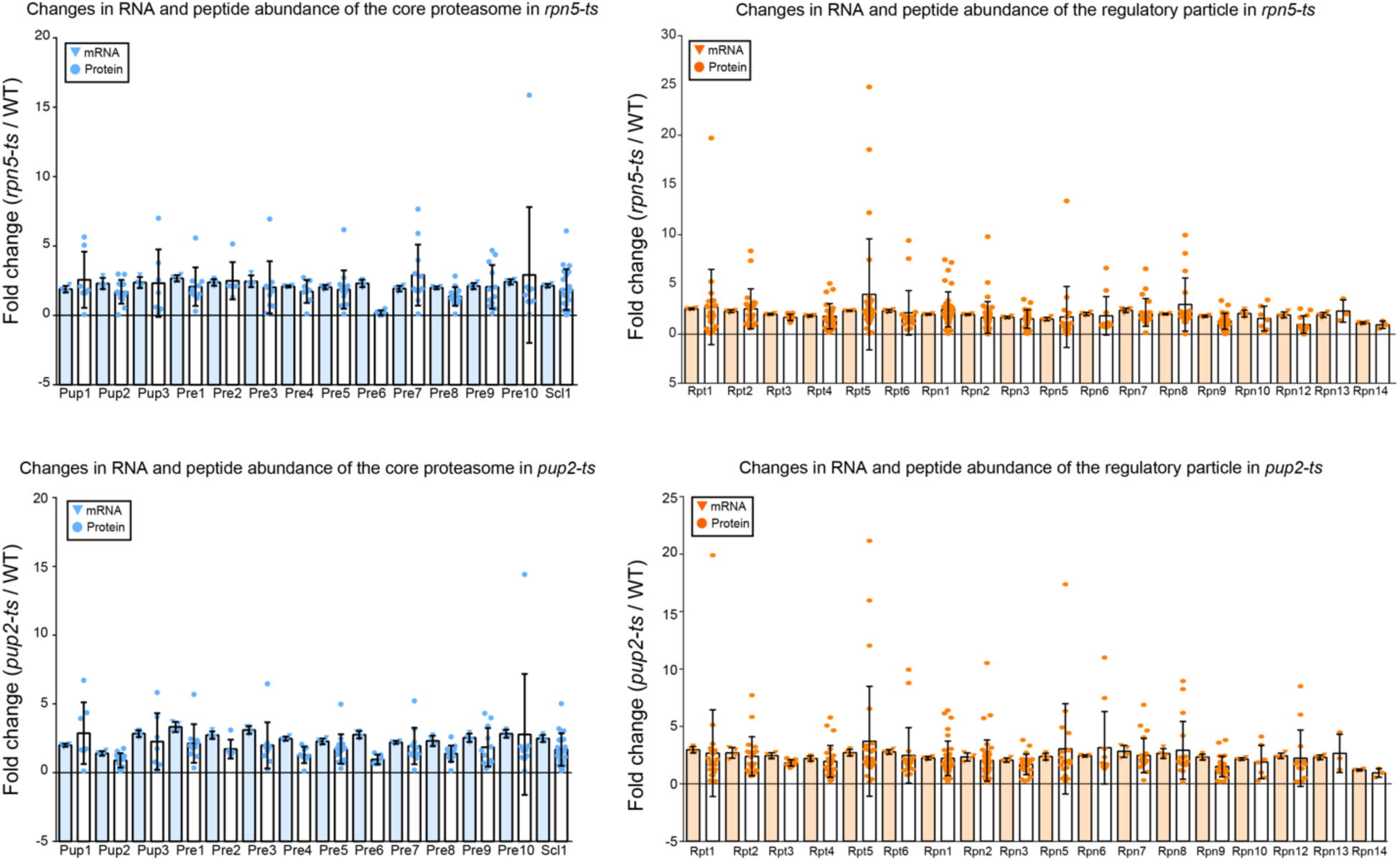
Comparing RNA and protein abundance for the subunits of the proteasome. Dot plots of protein and mRNA fold change in mutant/WT for each of the proteasome core (blue) and regulatory particle (orange) subunits. Shown are the unique peptide level fold changes (showing the average abundance value for all peptide-spectrum match for each peptide) from TMT-based quantitative global proteomics (n=3) and individual replicate fold changes from RNA sequencing experiments (n=4).

**Supplemental Figure 6:**
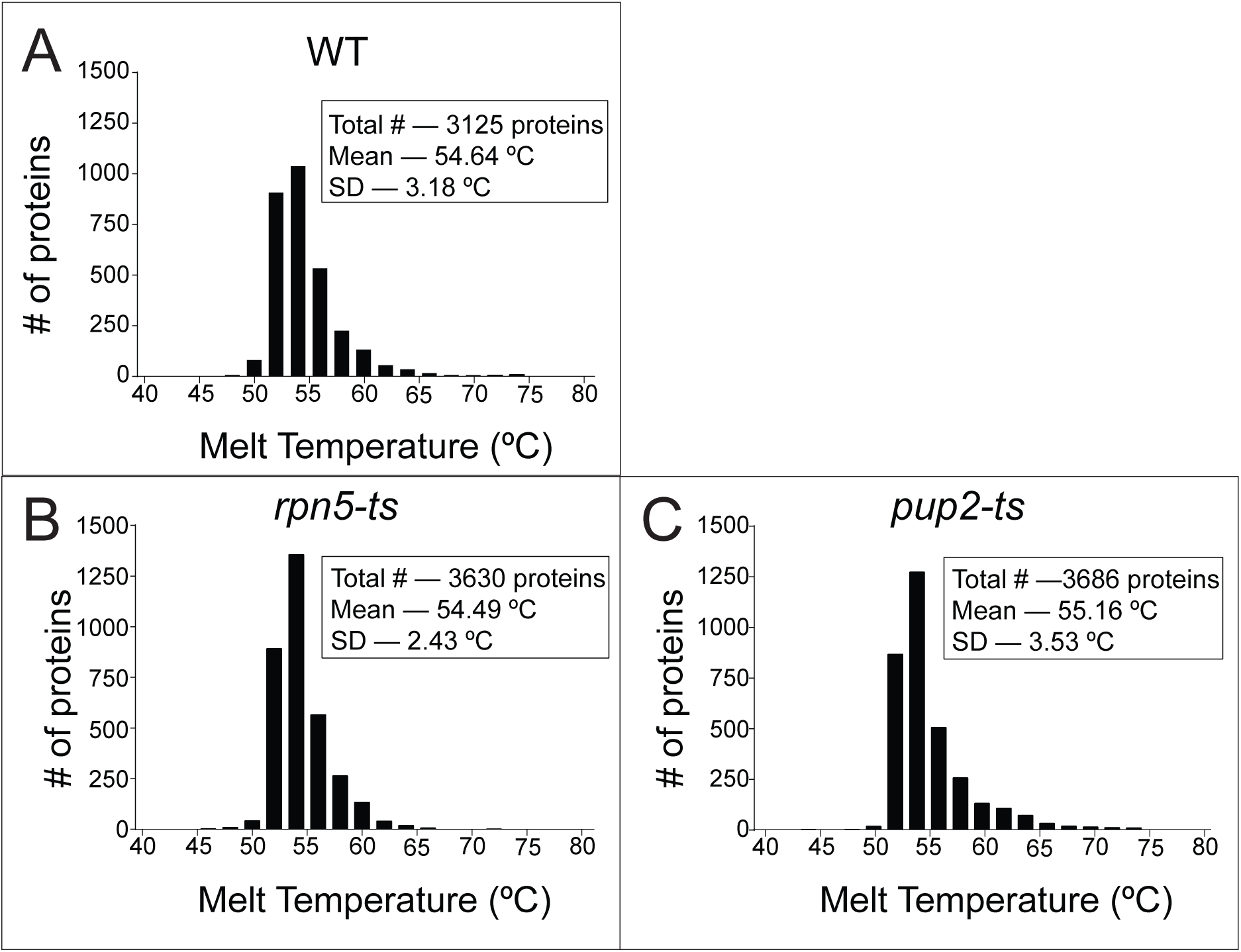
Distribution of protein melt temperature (T_m_) in WT, *rpn5-ts,* and *pup2-ts.* Shown is average T_m_ from TeMPP experiments of three biological replicates per genotype. Overall, the majority of proteins had a T_m_ within the 50-56 °C range. Plots were created and statistics were calculated using GraphPad Prism 6. A) A total of 2979 proteins were detected in WT. Average global T_m_ was 54.65°C. B) A total of 3284 proteins were detected in *rpn5-ts*. Average global T_m_ was 54.49°C. C) A total of 3297 proteins were detected in *pup2-ts*. Average global T_m_ was 55.16°C.

**Supplemental Figure 7 (part A):**
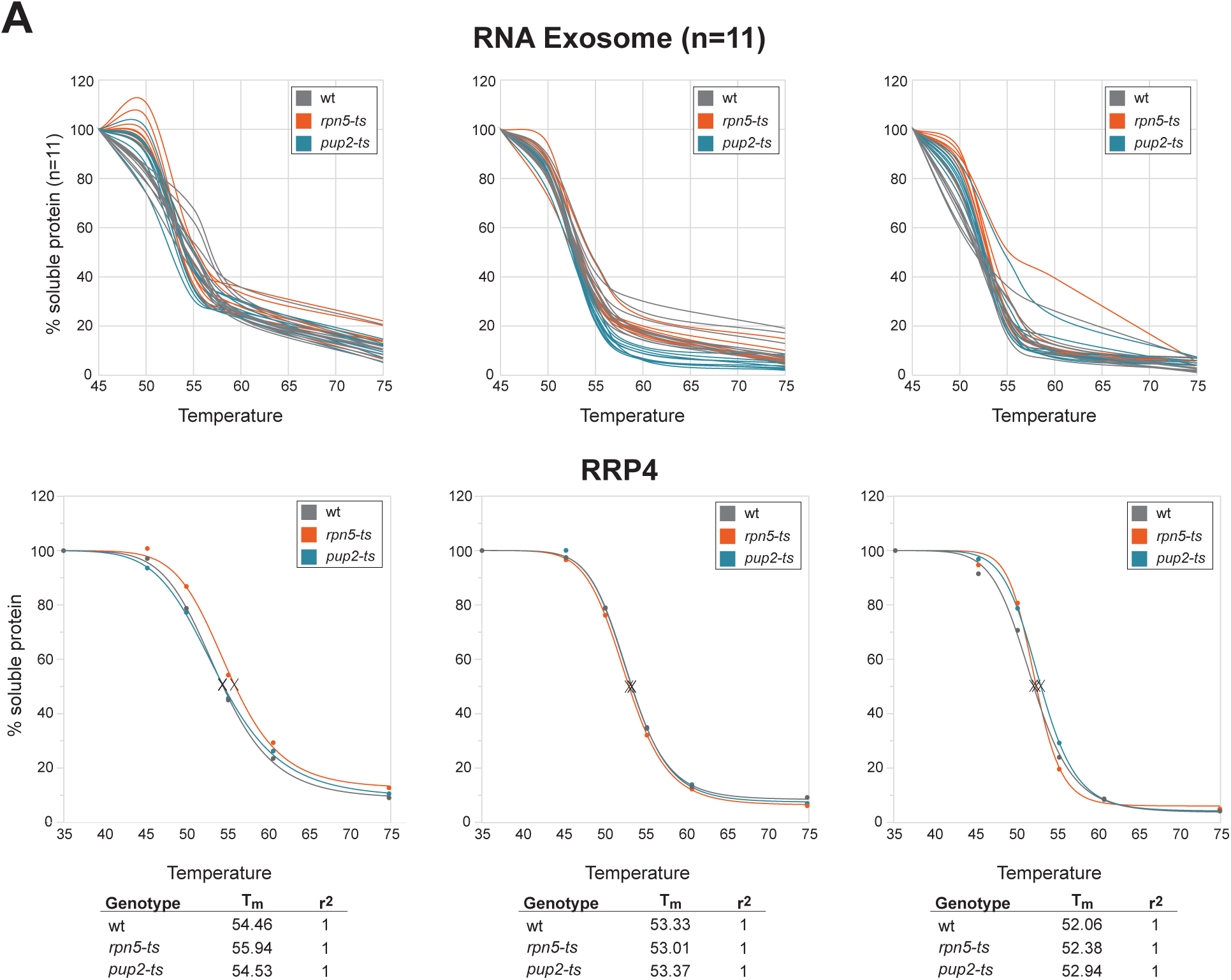
Replicates of complex melt curves and individual protein melt curves shown in Figure 1. Replicates of the melt curves shown in Figure 1 of the main text. Representative curves in the main text were all from the 2^nd^ biological replicate. Proteins isolated from wildtype are shown in gray, *rpn5-ts* in orange, and *pup2-ts* in teal. Individual protein melt curves were created for every quantified protein and normalized using the TPP R package. T_m_ was calculated as the temperature at which 50% of the protein was denatured. A) Replicate curves of the RNA exosome (11 individual proteins shown) and individual normalized curves from TPP of the exosome subunit Rrp4.

**Supplemental Figure 7 (part B):**
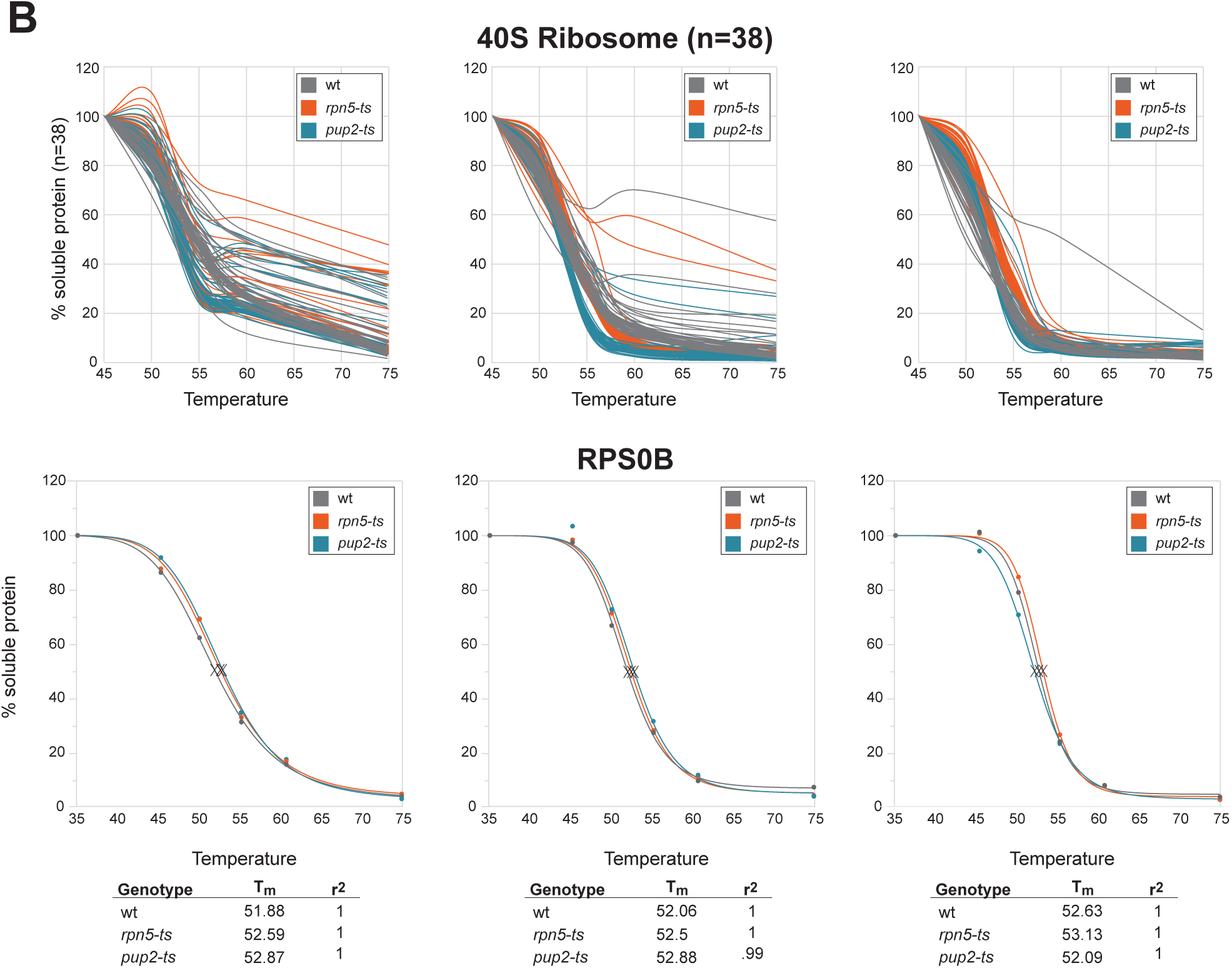
Replicates of complex melt curves and individual protein melt curves shown in Figure 1. Replicates of the melt curves shown in Figure 1 of the main text. Representative curves in the main text were all from the 2^nd^ biological replicate. Proteins isolated from wildtype are shown in gray, *rpn5-ts* in orange, and *pup2-ts* in teal. Individual protein melt curves were created for every quantified protein and normalized using the TPP R package. T_m_ was calculated as the temperature at which 50% of the protein was denatured. B) Curves of the 40S ribosome (39 individual proteins shown) and individual normalized curves from TPP of the ribosome subunit Rps0B.

**Supplemental Figure 7 (part C):**
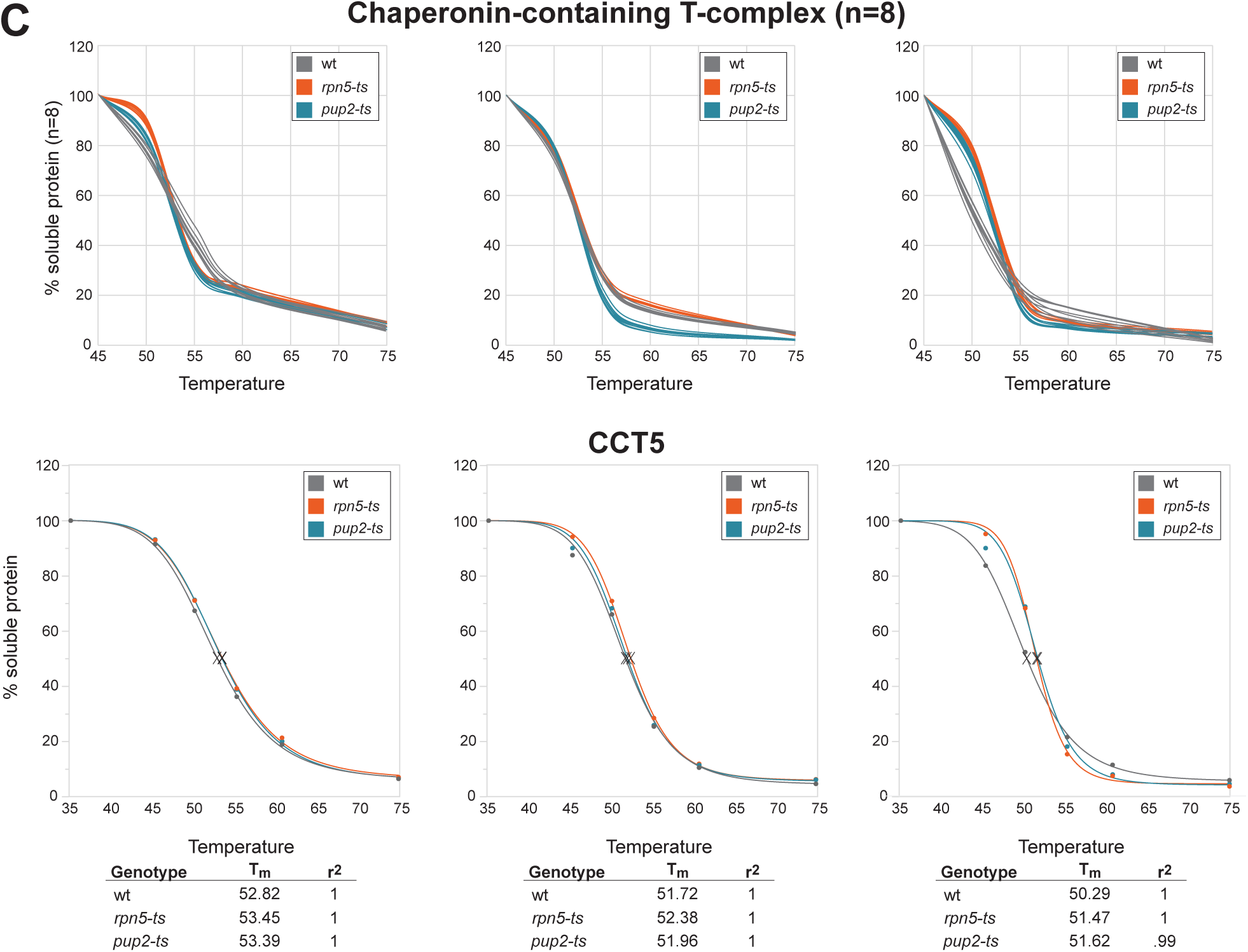
Replicates of complex melt curves and individual protein melt curves shown in Figure 1. Replicates of the melt curves shown in Figure 1 of the main text. Representative curves in the main text were all from the 2^nd^ biological replicate. Proteins isolated from wildtype are shown in gray, *rpn5-ts* in orange, and *pup2-ts* in teal. Individual protein melt curves were created for every quantified protein and normalized using the TPP R package. T_m_ was calculated as the temperature at which 50% of the protein was denatured. C) Curves of the Chaperonin-containing T-complex (8 individual proteins shown) and individual normalized curves from TPP of the CCT subunit Cct5.

**Supplemental Figure 8 (part A):**
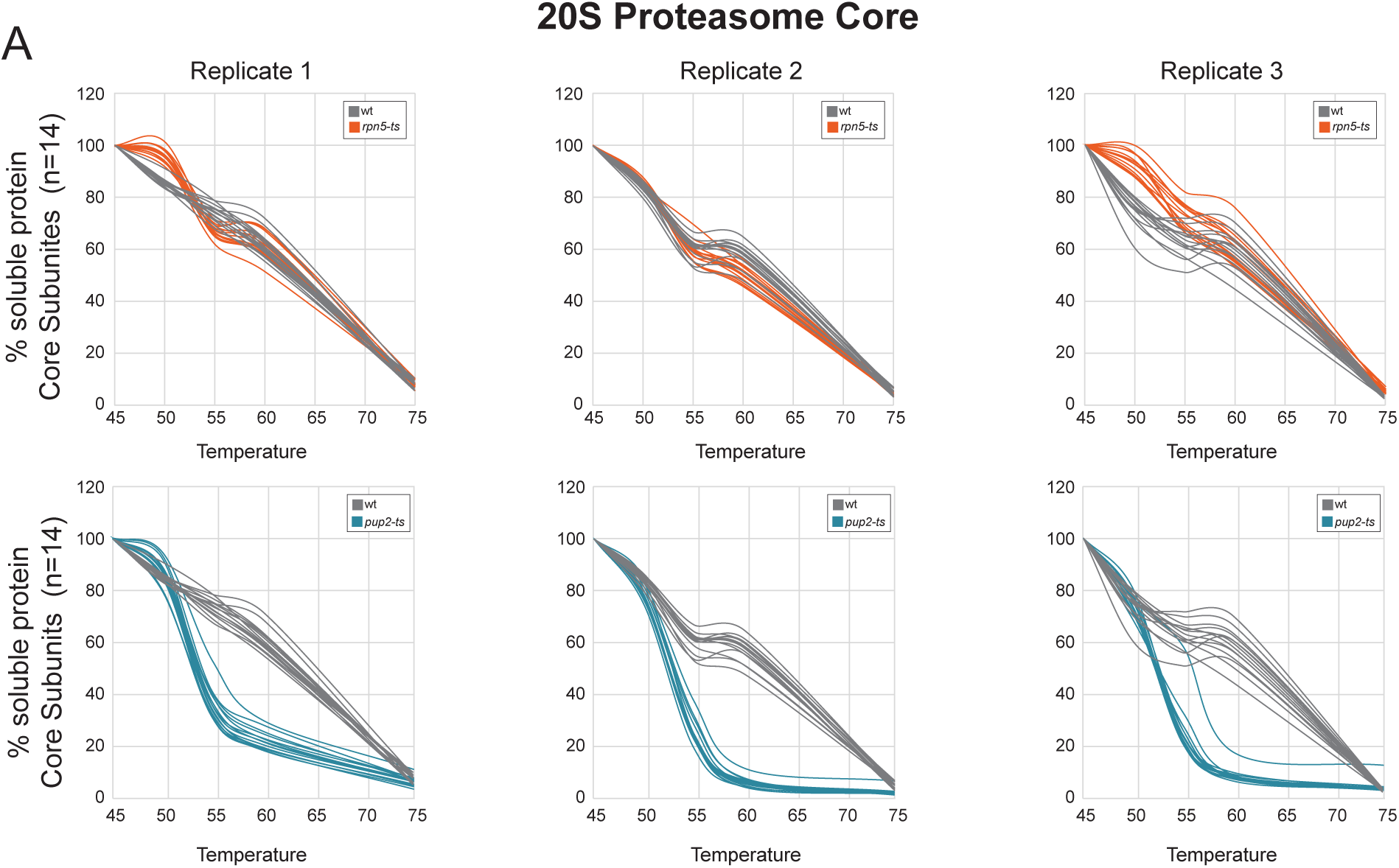
Proteasome melt curve replicates. Replicates of the proteasome melt curves shown in Figures 2&3 of the main text. Representative curves in the main text were all from the 2^nd^ biological replicate. A) Three replicates of curves of each of the 14 subunits of the 20S proteasome core in WT (gray) vs *rpn5-ts* (orange) or *pup2-ts* (teal). B) Three replicates of each curves of each of the 19 subunits of the 19S regulatory particle in WT vs. *rpn5-ts* (orange) or *pup2-ts* (teal).

**Supplemental Figure 8 (part B):**
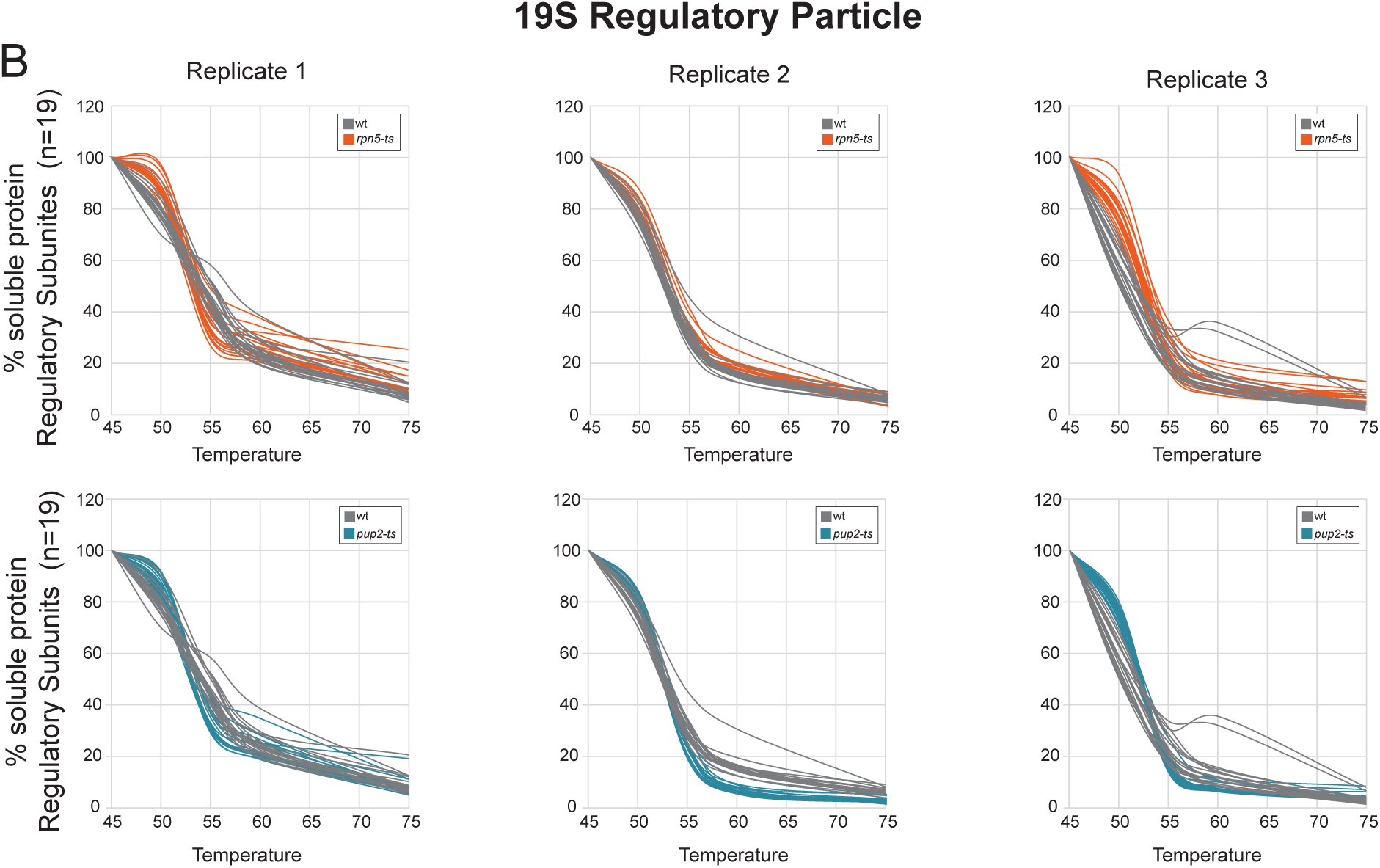
Proteasome melt curve replicates. Replicates of the proteasome melt curves shown in Figures 2&3 of the main text. Representative curves in the main text were all from the 2^nd^ biological replicate. B) Three replicates of each curves of each of the 19 subunits of the 19S regulatory particle in WT vs. *rpn5-ts* (orange) or *pup2-ts* (teal).

**Supplemental Figure 9:**
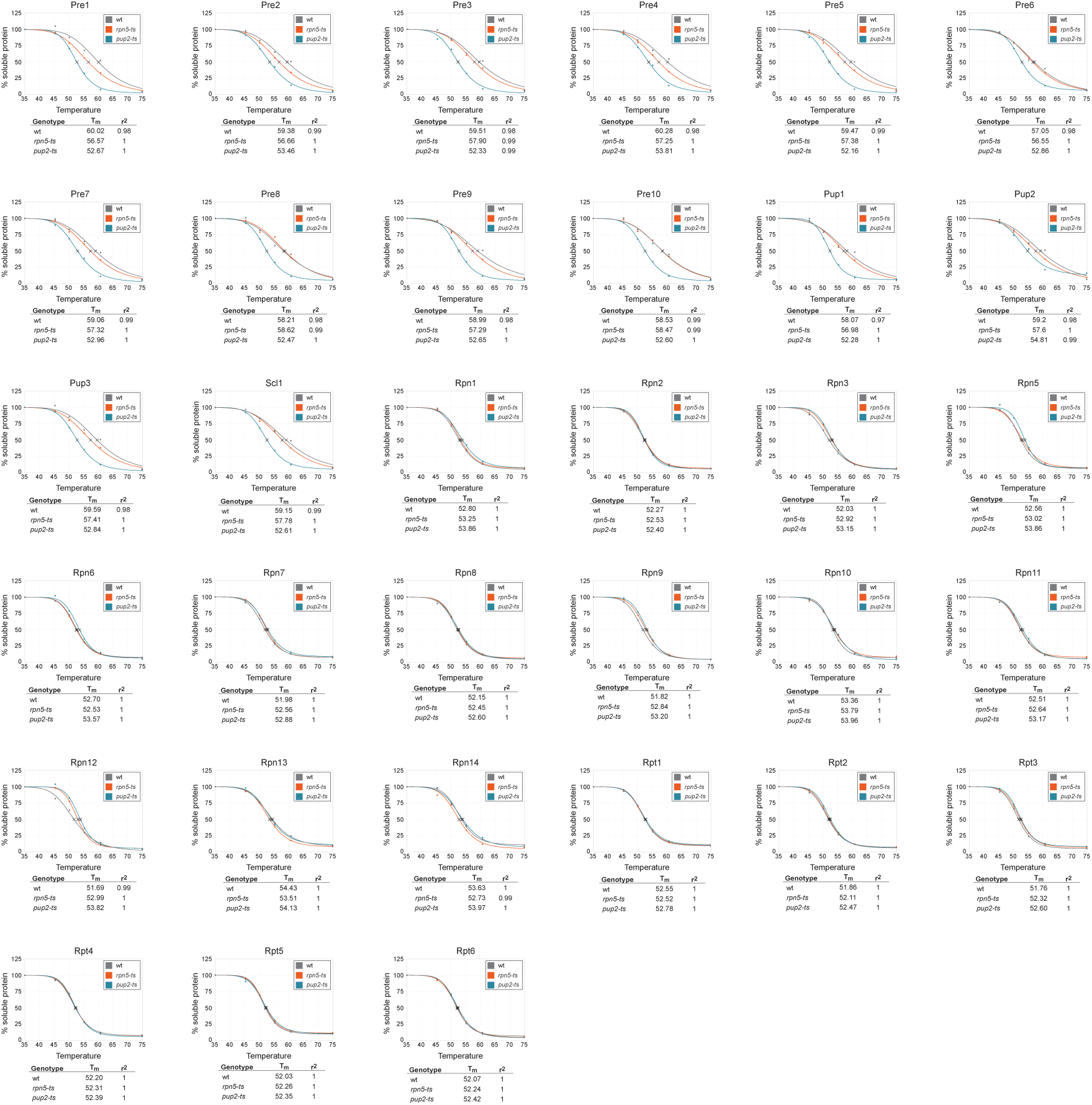
Normalized melt curves of the proteasome subunits from TPP. Plots for each of the individual proteasome subunits normalized by the TPP package for a representative replicate (p2). Curves shown in gray are WT, orange are from *rpn5-ts,* and blue are from *pup2-ts*.

**Supplementary Table 1: Global protein abundance data** Table of global proteomics data exported from Proteome Discoverer. Provided is the accession number, protein description from Uniprot, Sum PEP score, percent coverage, number of peptides, number of peptide spectral matches (PSMs), number of unique peptides, number of amino acids, molecular weight, abundance ratios, abundance ration p-values, grouped abundances for each genotype, normalized abundances for each TMT channel, and raw abundances for each TMT channel.

**Supplementary Table 2: RNA sequencing data** Table of RNA sequencing data for protein coding genes in WT, *pup2-ts,* and *rpn5-ts.* Provided is the gene name, Uniprot accession number, gene location information, fold change data and statistics, log_2_RPKM, and raw sequencing counts for each of the four replicates per genotype.

**Supplementary Table 3: TeMPP data** Table of data obtained from TeMPP experiments exported from Proteome Discoverer. Each genotype and replicate are provided as a separate sheet within the document. Provided is the accession number, protein description from Uniprot, Sum PEP score, percent coverage, number of peptides, number of peptide spectral matches (PSMs), number of unique peptides, number of amino acids, molecular weight, raw abundances for each channel, and percent of soluble protein as normalized to the lowest heat-treated sample (35°).

**Supplementary Table 4: TPP package results** Output results provided from the TPP package of the TeMPP data in Table 1.

**Supplementary Table 5: Changes in T_m_**Table of change in T_m_ (calculated from the results in Table 2) for each protein quantified in both mutant and WT. T_m_ values for *rpn5-ts* and *pup2-ts* were subtracted from WT to get changes in T_m_.

**Supplementary Table 6: Changes in T_m_ medians** Table of median values for change in T_m_. Median values were calculated for each protein that was quantified and had was given an T_m_ by the TPP package in at least two replicates.

**Supplementary Table 7: Proteins that have significant changes in thermal stability** Table of the proteins that significantly increased or decreased (using a 2σ cutoff) in thermal stability in *rpn5-ts* and *pup2-ts*.

